# Synaptic proteome diversity is shaped by the levels of glutamate receptors and their regulatory proteins

**DOI:** 10.1101/2024.04.04.588090

**Authors:** Rita Reig-Viader, Diego del Castillo-Berges, Albert Burgas-Pau, Daniel Arco-Alonso, Oriana Zerpa-Rios, David Ramos-Vicente, Javier Picañol, Aida Castellanos, David Soto, Nerea Roher, Carlos Sindreu, Àlex Bayés

**Affiliations:** Molecular Physiology of the Synapse Laboratory, Institut de Recerca Sant Pau (IR Sant Pau), Barcelona, Spain; Universitat Autònoma de Barcelona (UAB), Bellaterra (Cerdanyola del Vallès), Spain; Unitat mixta d’Investigació IRTA-UAB en Sanitat Animal. Centre de Recerca en Sanitat Animal (CReSA), Bellaterra (Cerdanyola del Vallès), Spain; IRTA. Programa de Sanitat Animal. Centre de Recerca en Sanitat Animal (CreSA), Bellaterra, 08193, Catalonia. Spain; Institute of Biotechnology and Biomedicine (IBB) and Department of Cell Biology, Physiology and Immunology, Universitat Autònoma de Barcelona, Bellaterra (Cerdanyola del Vallès), Spain; Neurodegenerative Diseases Research Group, Vall d’Hebron Research Institute, Centre for Networked Biomedical Research on Neurodegenerative Diseases (CIBERNED), Barcelona, Catalonia, Spain; Neurophysiology Laboratory, Department of Biomedicine, Faculty of Medicine and Health Sciences, Institute of Neurosciences, University of Barcelona, 08036 Barcelona, Spain; CIBER de Bioingeniería, Biomateriales y Nanomedicina (CIBER-BBN), Madrid, Spain; Institut d’Investigacions Biomèdiques August Pi i Sunyer (IDIBAPS), Barcelona, Spain and CIBER on Rare Diseases CIBERER, Instituto de Salud Carlos III

**Keywords:** Synaptic type, proteomics, proteome diversity, transcriptomics, laser-capture microdissection, hippocampus, trisynaptic circuit, glutamate receptors, gene regulation

## Abstract

Synapses formed by equivalent pairs of pre- and postsynaptic neurons have similar electrophysiological characteristics, belonging to the same type. However, these are generally confined to microscopic brain regions, precluding their proteomic analysis. This fact has greatly limited our ability to investigate the molecular basis of synaptic physiology. We introduce a procedure to characterise the synaptic proteome of microscopic brain regions and explore the molecular diversity among the synapses forming the trisynaptic circuit in the hippocampus. While we observe a remarkable proteomic diversity among these synapses, we also report that proteins involved in the regulation of the function of glutamate receptors are differentially expressed in all of them. Moreover, neuron-specific gene expression programs would contribute to their regulation. Here, we introduce a combined proteomics and transcriptomics analysis uncovering a previously unrecognised neuron-specific control of synaptic proteome diversity, directed towards the regulation of glutamate receptors and their regulatory proteins.

## Introduction

Proteomics research performed on synaptic biochemical preparations has established a very comprehensive catalogue of proteins with a synaptic function^1–7^. This central advance in brain research has nevertheless been limited by the requirements of biochemical fractionation procedures and the sensitivity of proteomics methods, which involve relatively large brain areas, such as the hippocampus or neocortex^6,8–11^. Yet, these brain samples are not homogenous, containing many different synaptic types^12^. Accordingly, proteomics research uncovers the composition of the average, or the prototypical, synapse in a given sample. However, to understand the molecular mechanisms orchestrating the functional states that a synapse can take, it is imperative to investigate individual synaptic types. This is arguably the most important technical hurdle to precisely elucidate the molecular mechanisms behind synaptic function, with implications on information processing, cognition and disease.

Synaptic types can be defined in different ways, for instance they can be chemical or electrical. They can also be defined based on their neurotransmitter, the pair of neurons forming them or as recently shown, according to the expression patterns of key scaffolding molecules^13,14^. In the present work a synaptic type refers to that formed by a specific pair of pre- and post-synaptic neurons. This is because there is an extensive electrophysiological literature showing that synapses defined by connectivity have different functional properties^12,15–18^. A paradigmatic example is to be found in the hippocampus, where functional differences between CA3-CA1 and DG-CA3 glutamatergic synapses are prominent^17^.

Several methodological approaches have appeared in recent years to get closer to the final goal of isolating individual synaptic types or even individual synapses. All of them have been performed in mice and rely on genomic manipulations. Some of these approaches used fluorescently tagged proteins to sort synaptosomal preparations^19–23^. These methods have allowed investigation of glutamatergic neurons in large brain regions, or to investigate the cell-surface proteome of mossy fibre synapses in CA3^21^. Other approaches took advantage of proximity labelling methods to define the proteome of inhibitory synapses or the synaptic cleft^24–26^. More recently, confocal imaging studies in mice expressing proteins of the Psd95 family tagged with different fluorophores, provided a glimpse at the daunting molecular diversity that excitatory synapses could have, without losing anatomical information^14,27^. These cutting-edge studies are starting to uncover a molecular diversity among synapses that could only be suspected until now. Nevertheless, these approaches are not fit to explore the large proteomic landscapes of local synaptic types, and have low translational power, as they cannot be used in human samples.

To address the molecular diversity between types of glutamatergic synapses, we leveraged on the topographical organization of the hippocampus, which contains one of the best studied neuronal circuits in the brain, the trisynaptic circuit. Synapses in this circuit are anatomically segregated, being found in three distinct hippocampal layers^28^. The first synapse localizes to the molecular layer of the dentate gyrus and is made between the axons of layer II neurons from the entorhinal cortex and the dendritic spines of the granule cells in the dentate (EC-DG synapse). The second synapse is found in the stratum lucidum of the CA3 subfield, made between granule cells and pyramidal neurons (DG-CA3 synapse). These synapses are structurally unique, as they are formed by large presynaptic boutons, the mossy fibre boutons, that contact equally big dendritic structures called thorny excrescences^17^. Finally, the third synapse is in the stratum radiatum of the CA1 subfield, made by axons leaving CA3 neurons and contacting the proximal dendrites of CA1 pyramidal neurons (CA3-CA1 synapse).

Importantly, within its corresponding layer, each one of these synapses accounts for the vast majority of all synapses therein. More specifically, studies on the numbers and types of neurons and synapses in these hippocampal layers indicate that over 90% of the synapses in the stratum radiatum of the CA1 subfield^29–32^ and the stratum lucidum^33–37^ correspond to CA3-CA1 and DG-CA3 synapses, respectively. Similarly, electron microscopy studies in the molecular layer have shown that 86-90% of all synapses correspond to those stablished between layer II entorhinal cortex excitatory neurons and granule cells^38^.

In this work we introduce a high-yield procedure that allows to characterise the proteomic diversity between glutamatergic synapses. This method has allowed us to investigate the molecular diversity of the synapses that form the trisynaptic circuit of the dorsal hippocampus. We also investigated expression of genes coding for synaptic proteins in 55 neuronal types from the hippocampus and subiculum. Together, our proteomics and transcriptomics analysis indicate that abundance differences in glutamate receptors and the proteins that regulate them are common drivers of proteome variability across synaptic types and that neuron-specific gene expression mechanisms participate in this regulation.

## Results

### Isolation of synaptic proteins from microscopic brain regions

To increase the anatomical resolution of synapse proteomics we have developed a procedure to extract synaptic proteins from microscopic brain regions. This method combines laser-capture microdissection (LCM) with enhanced extraction and recovery of synaptic proteins. We applied this procedure to perform deep proteomic profiling of the synaptic types constituting the trisynaptic circuit from the dorsal hippocampus.

In this procedure forebrains are dissected and rapidly snap-frozen prior to cryo-sectioning. Brains cannot be chemically fixed, as this negatively interferes with later proteomic analysis. We stablished maximum section thickness for effective LCM cutting to be 10 μm. Microdissection was performed in coronal slices encompassing the first 500 μm of the dorsal hippocampus (Suppl. Fig. 1a). As the pyramidal and granular layers, containing cell bodies, can be visually distinguished (Fig. 1a), they can be excluded, collecting only the synaptic-rich neuropile (Fig.1b-c). By dissecting fragments of 100 μm in width it is possible to have control over the hippocampal layer acquired (Suppl. Fig. 1b-c). From the dentate gyrus we obtained the Molecular Layer (ML, Fig. 1d), from CA3 we dissected the Stratum Lucidum (SL, Fig. 1e) and from CA1 the Stratum Radiatum (SR, Fig. 1c). The characteristic translucidity of the SL helped in localizing and collecting this layer. As the total area of the anterior hippocampus in a coronal section is around 2.3mm^2^ (^39^), we estimate that the tissue collected from CA1-SR, CA3-SL and dDG-ML corresponds to 8%, 6% and 7% of the entire hippocampus, respectively.

**Figure 1.**
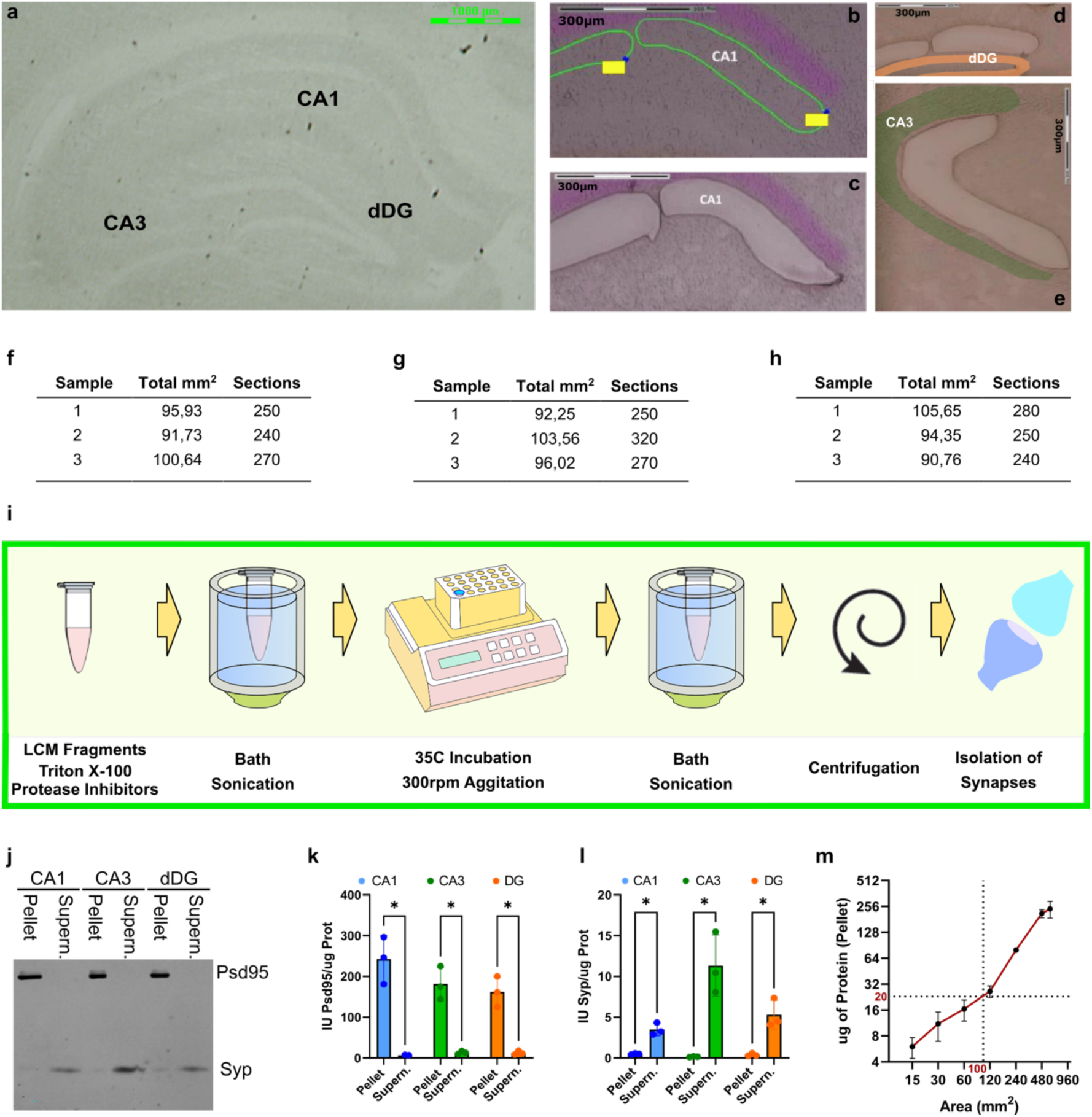
Effective separation of proteins from the synapses constituting the trisynaptic circuit using laser-capture microdissection and biochemical processing of hippocampal layers. **a.** Brightfield image of a coronal brain section showing the hippocampus, as used for laser-capture microdissection (LCM). Note that CA1/CA3 pyramidal layers and dDG granular layer are visible. Scale bar 1000μm. **b.** CA1 subfield before microdissection. Pyramidal Layer in purple. Green line marks microdissected area. Microdissected fragments had a width of approximately 100μm, only collecting neuropile from the Stratum Radiatum. Scale bar 300μm. **c.** CA1 subfield from the section shown in (b) after LCM. The pyramidal layer, in purple, is not collected. Scale bar 300μm. **d.** Dentate gyrus after LCM. Microdissected fragments had a width of approximately 100μm, which allowed specifically collecting neuropile from the Molecular Layer. Granular layer highlighted in orange. Scale bar 300μm. **e.** CA3 subfield after LCM. Microdissected fragments had a width of approximately 100μm, which allowed to collect neuropile from the Stratum Lucidum. Pyramidal layer in green. Scale bar 300μm. **f.** Total area (mm^2^) microdissected and number of brain sections collected for the biological replicas used in proteomics of dDG. **g.** Total area (mm^2^) microdissected and number of brain sections collected for the biological replicas used in proteomics of CA3. **h.** Total area (mm^2^) microdissected and number of brain sections collected for the biological replicas used in proteomics of CA1. **i.** Outline of the procedure used to enrich LCM samples in synaptic proteins. **j.** Immunoblot of 1% Triton X-100 insoluble (Pellet) and soluble (Supern.) fractions from the three hippocampal layers investigated. Proteins analysed are Psd95 and Synaptophysin (Syp). **k.** Relative Psd95 abundance determined by immunoblot in 1% Triton X-100 soluble (Supern.) and insoluble (Pellet) fractions. IU: intensity units. Error bars: SE. Sample size (n) = 3. Statistics, Two-way ANOVA and Fisher’s LSD post-hoc test, * p < 0.05. **l.** Relative Synaptophysin abundance (Syp) determined by immunoblot in 1% Triton X-100 soluble (Supern.) and insoluble (Pellet) fractions. IU: intensity units. Error bars: SE. Sample size (n) = 3. Statistics, Two-way ANOVA and Fisher’s LSD post-hoc test, * p < 0.05. **m.** Micrograms of protein recovered in 1% Triton X-100 pellets per area of microdissected neuropile. To obtain 20μg of protein 100mm^2^ of neuropile have to be microdissected. Error bars: SE.

Extracting synaptic proteins from the microscopic amounts of tissue collected by LCM is very challenging. To cope with this limitation, we developed a procedure designed to minimize sample manipulation, reducing sample loss, while maximizing recovery of synaptic proteins. This procedure takes advantage of the selective solubility of synaptic structures to the detergent Triton X-100, such as the postsynaptic density (PSD), the active zone (AZ) or the extracellular matrix of the synaptic cleft^5^. First, microdissected tissue is accumulated in a solution containing 1% Triton X-100 (Fig. 1f-h). Next, neuropile fragments are subjected to a three-step treatment, a brief bath sonication, a mild thermal shock at 35C in agitation, and a second sonication step. This procedure fully disperses neuropile fragments and maximises the effect of the detergent, while preserving protein integrity and avoiding sample manipulation. A final centrifugation allows to collect Triton-insoluble proteins (Fig. 1i).

To evaluate the efficacy of this procedure, we assayed samples by immunoblot against proteins known to be mostly soluble (Synaptophysin, Syp) or mostly insoluble (Psd95) to Triton X-100. Over 90% of the Psd95 signal was detected in pellets, conversely, the same proportion of Syp was in supernatants (Fig.1j-l). No difference in Psd95 abundance was observed between samples (two-way ANOVA), indicating that the procedure had a similar efficiency in all hippocampal layers.

As these samples contain very little protein, standard approaches for protein quantification could not be used. Protein concentration was determined by electrophoresis, using as internal calibration standards hippocampal synaptic preparations accurately quantified (Suppl. Fig. 2a,b). Using this approach, we determined that insoluble fractions contain approximately 20% of all protein in the tissue (Suppl. Fig. 2c), indicating that proteins in these fractions were concentrated 4-5 times. We also tested different extraction buffers to investigate if we could improve the efficiency of the procedure. Using a RIPA buffer we found that the amount of protein recovered in pellets was significantly smaller (Suppl. Fig. 2d,e), yet this was at the expense of solubilizing a larger proportion of both Psd95 and Syp (Suppl. Fig. 2f-g). Indicating that more synaptic components were lost in the soluble fraction. On the other hand, increasing Triton concentration to 2% did not improved protein yield (Suppl. Fig. 2e). Neither RIPA nor 2% Triton showed improved performance over 1% Triton X-100, which remained as the buffer of choice. Finally, we established how much protein was recovered in pellets per area of microdissected neuropile, this was important to keep LCM time to a minimum. We determined that for each 100mm^2^ of neuropile we obtained approximately 20μg of Triton insoluble protein (Fig. 1m).

### High similarity in the composition of trisynaptic circuit synapses

Using the above procedure, we obtained biological triplicates of synaptic preparations from the layers of the trisynaptic circuit and subjected them to a proteomics workflow^40^. MS/MS data was examined with Scaffold-DIA (Proteome Software), to identify protein specimens, and Progenesis QI (Waters), for peptide quantification (Fig. 2a). Peptide abundance was normalized by the average abundance of peptides from 14 synaptic scaffolding proteins (see methods). This allowed us to correct for differences in: i) synaptic yield between preparations and ii) synaptic density between layers. Finally, MsqROB^41,42^ was used to identify proteins differentially expressed (DE) between synaptic types.

**Figure 2.**
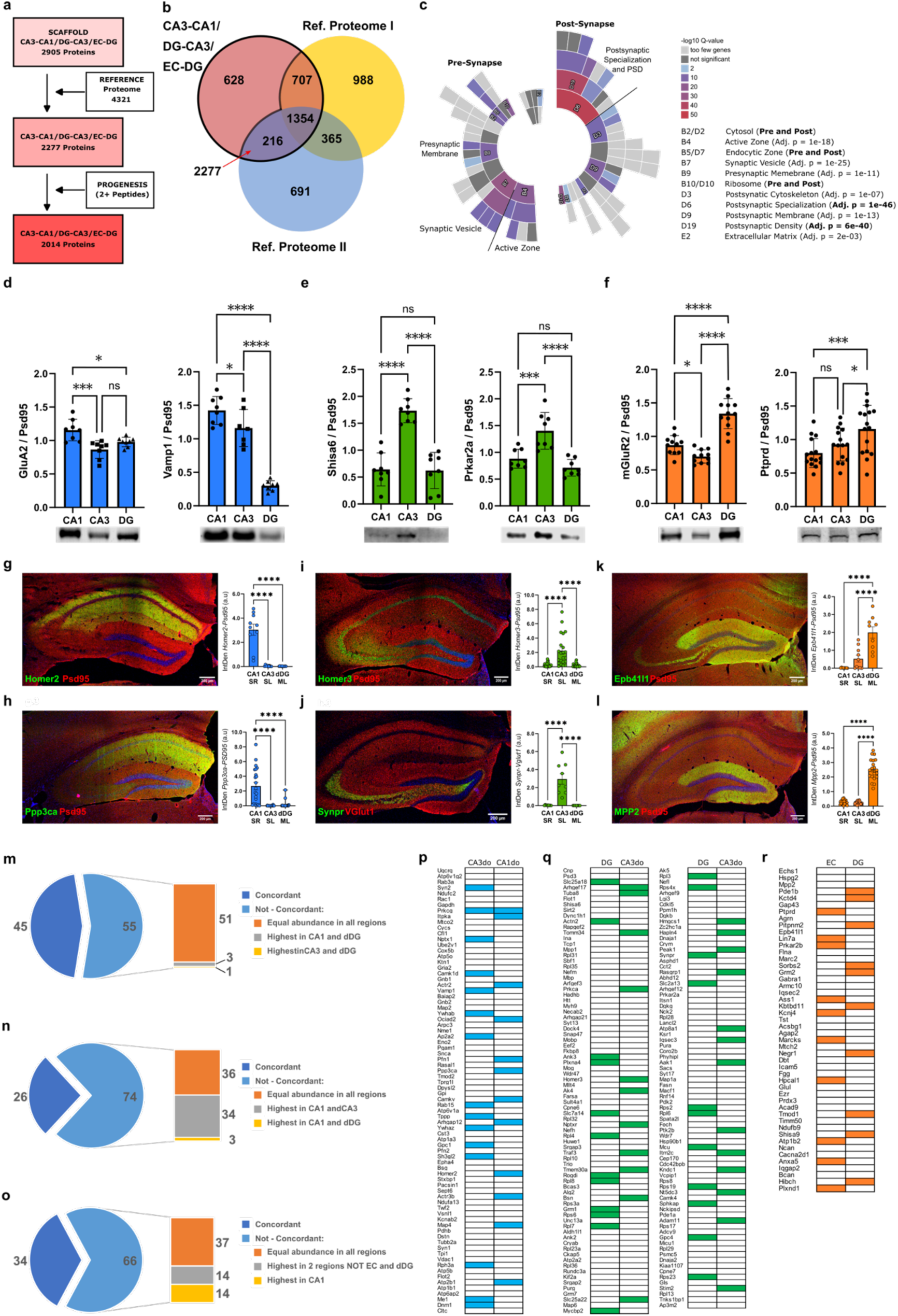
Proteomics workflow and validation of proteins found differentially expressed. **a.** Steps involved and informatic tools used to establish the final proteome of synapses from the trisynaptic loop. **b.** Overlap between proteins from the trisynaptic loop and two reference proteomes. Ref. Proteome I, from this study; Ref. Proteome II is PSDII from Distler et al^8^. **c.** Sunburst plot showing SynGO Cellular Component terms enriched among synaptic proteins from the trisynaptic loop. **d-f.** Relative abundances of GluA2 (d, left), Vamp1 (d, right), Shisa6 (e, left), Prkar2a (e, right), mGluR2 (f, left) and Ptprd (f, right) determined by immunoblot in synaptic fractions from CA1, CA3 and DG subfields. Representative immunoblots shown. Error bars: SE. Sample size (n) = 6, 2-3 replicates per sample. Statistical test, one-way ANOVA, post-hoc Fisher’s LSD test, * p < 0.05, *** p < 0.001, **** p < 0.0001. **g-l. Left:** Representative images of double immunofluorescences of the postsynaptic marker Psd95 (in red) with Homer2 (g), Calcineurin (Ppp3ca, h), Homer3 (i), Band 4.1-like protein 1 (Epb41l1, k) and Mpp2 (l) and the presynaptic marker vGlut1 (in red) with Synaptoporin (Synpr, j). **Right:** Quantification of the overlapping signal in IFs (see also Suppl. Figs. 4-6 for more detail on the IFs): Homer2 (g), Ppp3ca (h), Homer3 (i), Synpr (j), Epb41l1 (k) and Mpp2 (l). Error bars: SE. Sample size (n) = 4, 3-6 images taken from subfield and animal. Statistical test, one-way ANOVA, post-hoc Fisher’s LSD test, **** p < 0.0001. **m.** Percentage of DE proteins in CA3-CA1 synapses with concordant or discordant ISH levels. **n.** Percentage of DE in DG-CA3 synapses with concordant or discordant ISH levels. **o.** Percentage of DE proteins in EC-DG synapses with concordant or discordant ISH levels. **p.** DE proteins in CA3-CA1 synapses with increased scRNAseq levels in dorsal CA3 (CA3do, left column) or dorsal CA1 (CA1do, right column) neurons are indicated with a blue box. **q.** DE proteins in DG-CA3 synapses with increased scRNAseq levels in dentate gyrus (DG, left column) or dorsal CA3 (CA3do, right column) neurons are indicated with a green box. **r.** DE proteins in EC-DG synapses with increased scRNAseq levels in Entorhinal cortex (EC, left column) or Dentate Gyrus (DG, right column) neurons are indicated with an orange box.

The proteomic dataset obtained from microdissected tissue was benchmarked against a reference proteome. This was generated from the combination of two proteomes of hippocampal synaptic fractions prepared by differential ultracentrifugation (Suppl. Fig. 2i)^6^. We generated one of these datasets and the other had been previously published^8^ (Fig. 2b and Suppl. Table 1). Proteins detected in LCM samples but absent from the reference proteome were discarded as contaminants (Fig. 2a). An analysis of overrepresentation of synaptic locations (GO-CCs) and biological processes (GO-BPs) among discarded proteins returned only one significant GO-CC (postsynaptic ribosome) and no significant GO-BPs (Suppl. Fig. 3a,b). Indicating that benchmarking served the purpose of removing contaminants from our synaptic preparations. Of the 2905 identified by Scaffold 2277 remained after benchmarking. Of these, 2014 could be quantified with at least 2 peptides by Progenesis, this being the final dataset investigated (Fig. 2a and Suppl. Table 1).

We next confirmed that our method was able to retrieve proteins from distinct subsynaptic locations. Using the SynGO database^2^ to assign subsynaptic compartments onto our dataset, we found that it was enriched in many of them, both pre- and postsynaptic (Fig. 2c). As a matter of fact, pre- and postsynaptic proteins were similarly enriched. The presence of presynaptic proteins in our preparations was confirmed by immunoblot (Suppl. Fig. 2f,j). Thus, this approach provides a wide view into the synaptic proteome.

A small number of proteins were identified only in one sample (CA3-CA = 29, DG-CA3 = 68 and EC-DG = 52, Suppl. Table 1). Potentially these proteins could be very interesting, as they might be markers of synaptic types. Nevertheless, most of them (86%) could only be identified in one of the three replicates, and their abundance was very low (mean 3.45 peptides/protein, compared with 43 peptides/protein for the whole set). Thus, we decided to exclude these molecules from subsequent analysis. Our data suggests that few proteins, if any, are unique to a single synaptic type in the trisynaptic loop, making functionally different synapses nearly identical at the proteomic level.

### Identification and validation of differentially expressed synaptic proteins

The above data implied that quantitative, rather than qualitative variation drives functional diversity across synapses. To identify DE synaptic proteins, we used a ridge regression method designed to analyse peptide abundances acquired by label-free mass spectrometry^41,42^. We identified a total of 283 significantly overexpressed proteins, 14% of the entire dataset, of which 78, 157 and 48 in CA3-CA1, DG-CA3 and EC-DG synapses, respectively (Suppl. Fig. 3c and Suppl. Table 2).

To validate our proteomics results we first performed immunoblot analysis of six DE proteins (GluA2 and Vamp1 for CA1; Shisa6 and Prkar2a for CA3; mGluR2 and Ptprd for DG; Fig. 2d-f) in synaptic fractions obtained from manually dissected hippocampal fields (Suppl. Video 1), finding perfect agreement between proteomics and immunoblot data. Afterwards, we performed double immunofluorescence (IF) on hippocampal slices for another 6 DE proteins (Homer2 and Ppp3ca for CA1-CA3, Homer3 and Synpr for DG-CA3, and Epb41l1 and Mpp2 for EC-DG synapses) and a pre- or a postsynaptic marker, to specifically look at their synaptic abundances. For each double IF we quantified the intensity of the overlapping signal in CA1-SR, CA3-SL and dDG-ML. This was always highest in the synapse where proteomics also identified maximum abundance, corroborating our initial findings (Fig.2g-l, and see Suppl. Figs. 4-6 for more detail on this analysis).

Finally, we also performed an electrophysiological validation of our proteomics data. Based on the fact that GluA2 was more abundant in CA3-CA1 synapses (Suppl. Table 2, Fig. 2d and Fig. 3), and that GluA2-containing AMPARs present slower deactivation kinetics^43–45^ regardless of the auxiliary subunits interacting with them^46,47^, we investigated if the decay of miniature excitatory postsynaptic currents (mEPSCs, Suppl. Fig. 7a-d) was slower in CA1 than in CA3 pyramidal neurons. mEPSCs were not investigated in DG granule cells as GluA2 levels in DG and CA3 synapses were very similar (GluA2 abundance ratio DG:CA3 0.94, Suppl. Table 2). mEPSCs had similar amplitudes in CA1 and CA3 neurons (Suppl. Fig. 7e) but presented a significantly slower decay in CA1 neurons. With a weighted mean time constant (T_w_) of 6.82ms for CA1 and 2.48ms for CA3 neurons (Suppl. Fig. 7f,g). This different kinetics would agree with an increased number of GluA2-containig AMPARs in CA3-CA1 synapses, as indicated by the proteomics data.

**Figure 3.**
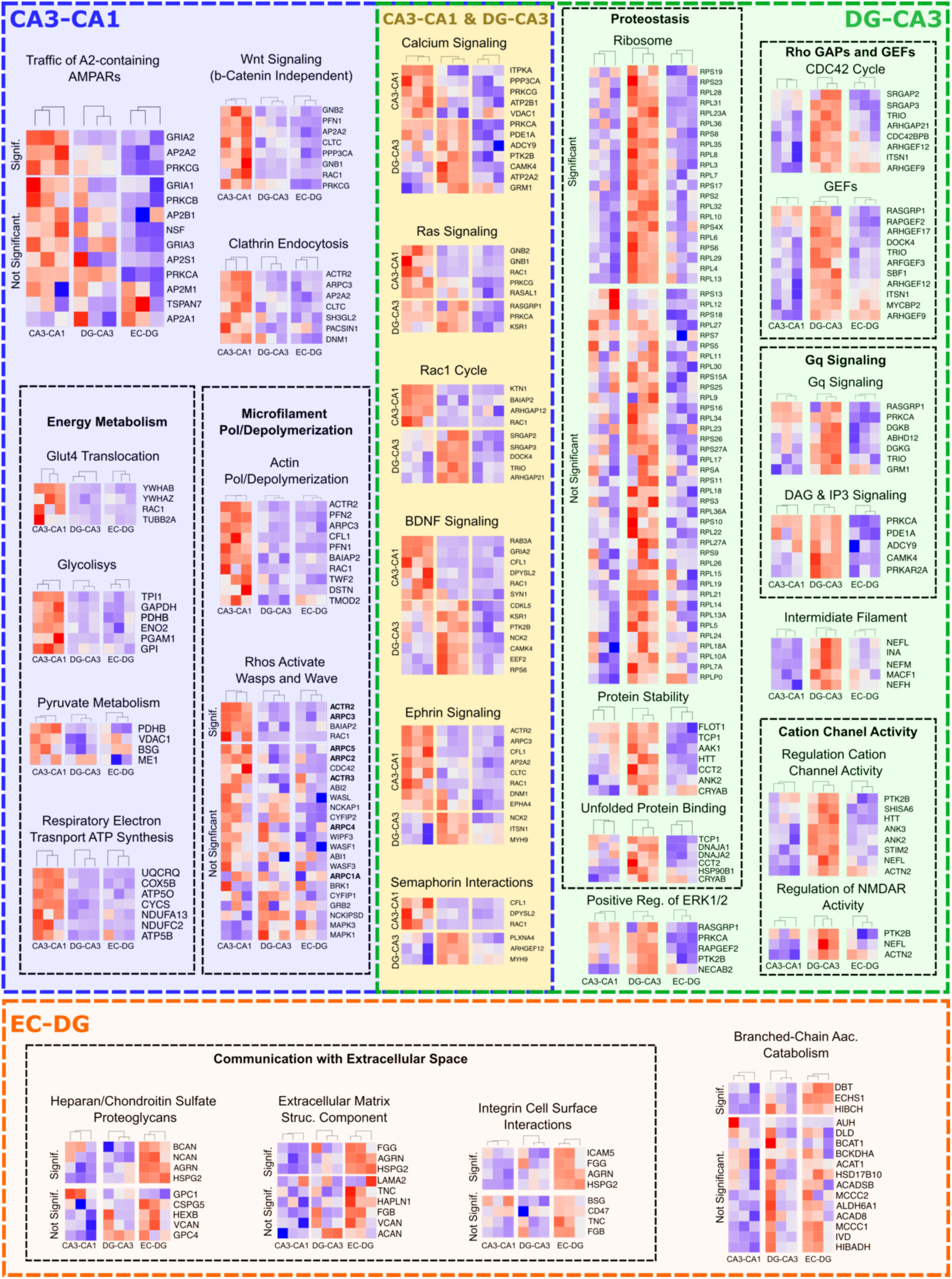
Signalling pathways differentially regulating function in each synaptic type from the trisynaptic circuit. Signalling pathways overrepresented amongst DE proteins in each synapse of the trisynaptic loop. Pathways specific to CA3-CA1 synapses are framed in a blue box, pathways specific to DG-CA3 in a green box, those common to these 2 in a yellow box and, finally, pathways specific to EC-DG synapses are in an orange box. Relative protein abundance for each of the 9 samples investigated by LC-MS/MS are presented as z-scores in heatmaps. A title and a heatmap is presented for each overrepresented pathway. Related pathways (i.e. CA3-CA1 pathways involved in Energy Metabolism) are framed with a dashed black line. For some pathways (i.e. Traffic of A2-containing AMPARs) we also present a heatmap with proteins that have a clear DE but did not reach statistical significance (Not Significant). In the ‘Rhos Activate Wasps and Waves’ gene names of members of the Arp2/3 complex are in bold.

### Contribution of gene expression to synaptic proteome diversity

Having identified and validated DE proteins across synaptic types, we asked if gene expression mechanisms could account for some of this variability. To achieve this, we analysed RNA expression data from *in situ* hybridization studies on the mouse brain (ISH, Allen Mouse Brain Atlas^48^, Suppl. Table 3) and single-cell RNA sequencing (scRNAseq) of excitatory neurons from the dorsal CA1, dorsal CA3, DG and entorhinal cortex (Allen Brain Cell Atlas, ABCA^49^, Suppl. Table 3), and compared them with proteomic abundances. Protein and RNA data were considered concordant if an upregulated protein showed highest RNA expression in the pre- and/or postsynaptic neuron/s forming it. We observed a protein to RNA abundance concordance of 35% and 34 % for ISH (Fig. 2m-o) and scRNAseq data (Fig. 2p-r, Suppl. Table 3), respectively. Importantly, a permutation test shows that this percentage of concordance is significantly higher than what would be expected by chance, as random concordance between our proteomics and the RNA expression data investigated would be around 23% (permutation test p=0.002, see Methods). What indicates that gene transcription partially accounts for synaptic proteome variability at the hippocampus.

### High diversity in the molecular mechanism operating at different synapses

To investigate the biological functions related to proteins with highest expression in one synaptic type, we performed enrichment analysis of signalling pathways^50–52^ and GO terms^53,54^ using the pathfindR tool^55^. pathfindR constructs protein-protein interaction networks and maps enriched terms onto them. Using hierarchical clustering and pairwise kappa statistics, pathfindR identifies one ‘Representative’ term for each network (see methods).

We first clarified if a small number of proteins were responsible for a large proportion of enriched terms, a common bias with pathway enrichment analysis^56,57^. Yet this was not the case, as the ratio of enriched terms per protein was low (Suppl. Fig. 3d) and the proportion of proteins contributing to terms was high (Suppl. Fig. 3e) in all samples. Importantly, most enriched pathways (75%) and GO terms (96%) were found only in one synaptic type (Suppl. Fig. 3f,g), thus, effectively informing about their unique functional properties. Only 5 terms were enriched in all samples. These were strongly related to synaptic function and included transmission across chemical synapses, postsynaptic signalling, actin cytoskeleton and cell adhesion (Suppl. Fig. 3h and Suppl. Table 4).

While CA3-CA1 and DG-CA3 synapses shared several functional categories, only 3 were found between DG-CA3 and EC-DG synapses and none between CA3-CA1 and EC-DG synapses. Of note, the GOCC term ‘Schaffer collateral CA1 synapse’, appeared enriched in DE proteins from CA3-CA1 and DG-CA3 synapses. Among the pathways common to these synapses we identified well-known synaptic processes, such as signalling via calcium, through Ras and Rho GTPases or trans-synaptic signalling via BDNF, Ephrins and Semaphorins (Fig. 3).

### AMPAR trafficking, actin dynamics and energy metabolism regulated in CA3-CA1 synapses

As previously mentioned, we observed an increased abundance of the GluA2 AMPAR subunit in these synapses (Fig. 2d and 3), suggesting that they would have more Gria2-containig AMPARs, an observation supported by our recordings of mEPSCs (Supp. Fig. 7). In agreement with these findings the analysis of enriched pathways retrieves ‘Traffic of GluA2 containing AMPAR’ as strongly overrepresented among DE proteins in these synapses (fold enrichment, 38.6, Suppl. Table 4). Other proteins involved in the regulation of AMPAR trafficking, such as those controlling clathrin-mediated endocytosis^58^ and neuronal pentraxin 1 (Nptx1)^59^, were also strongly enriched in these synapses.

Although actin-related categories were found in all synaptic types (Suppl. Fig. 3h and Suppl. Table 4), CA3-CA1 synapses presented many more functional categories related to microfilaments, particularly to their polymerization. For example, all 7 members of the Arp2/3 complex, necessary for actin branching and dendritic spine structural plasticity^60^, presented higher abundance in this synaptic type, albeit only three reached statistical significance (Fig. 3 and Suppl. Table 2). This would be suggestive of a more refined control of spine structural dynamics in these synapses.

We also found the non-canonical Wnt signalling pathway, which controls calcium levels and synaptic plasticity^61,62^ overrepresented in CA3-CA1 synapses. Among the downstream effectors of this pathway, calcineurin (Ppp3ca) and the calcium-activated protein kinase C (PKC, isoenzyme Prkcg) were overexpressed in this synaptic type, suggesting that the modulation of spine calcium dynamics via Wnt signalling might be especially relevant in these synapses.

Finally, multiple functional categories related to energy production were specifically overrepresented in CA3-CA1 synapses. Suggesting that these synapses would have higher energetic demands. These include proteins regulating the trafficking of glucose transporters to the plasma membrane, five out of the 10 glycolytic enzymes and enzymes related to pyruvate metabolism or ATP synthesis.

### Control over metabotropic signalling, translation and neurofilaments in DG-CA3 synapses

The postsynaptic metabotropic glutamate receptor Grm1 presented increased abundance in DG-CA3 synapses, particularly in relation to CA3-CA1 synapses, with a 3.4-fold increase. Grm1 signals through Gq protein alpha subunits, which regulate levels of the second messenger inositol trisphosphate (IP3) and diacyl glycerol (DAG). The signalling pathways ‘G alpha Q signalling events’ and ‘DAG and IP3 signalling’ were also found significantly enriched in DG-CA3 synapses. Similarly, Necab2 and Homer3, known to modulate metabotropic glutamate signalling^63^ were found strongly overexpressed in DG-CA3 synapses.

DE proteins in DG-CA3 synapses also regulate NMDA and AMPA receptors. We found overrepresented pathways related to NMDA receptor function, including ‘Regulation of NMDA Receptor Activity’ or ‘Negative Regulation of NMDA Receptor Mediated Neuronal Transmission’ (Suppl. Table 4). Among DE proteins controlling NMDARs, PTK2B might be particularly relevant, as this kinase also interacts with Grm1^64^. We also identified proteins regulating AMPAR function, including Shisa6^65^, Syt17^66^, Snap47^67^, and Nptxr^59^. Also related to the function of both AMPA and NMDA receptors is the signalling through ERK1/2 kinases. The GO pathway ‘Positive Regulation of ERK1 and ERK2 Cascade’ was also found overrepresented in DG-CA3 synapses.

Interestingly, among NMDAR related proteins we identified the neurofilament light chain (Nefl), known to be involved in its trafficking^68,69^. Actually, the four proteins that form neurofilaments were found significantly overexpressed in DG-CA3 synapses (Suppl. Table 2). Many modulators of the Rho family of small GTPases, including GTPase activating proteins (GAPs) and, specially, guanine nucleotide exchange factors (GEFs) were also found overexpressed. This suggests that pathways regulated by these signalling molecules, mostly related to the regulation of the cytoskeleton, might be controlled in a more specific manner in this synaptic type.

Finally, we observed a striking increase of virtually all ribosomal proteins in DG-CA3 synapses, with 21 of them reaching statistical significance (Fig. 3, Suppl. Tables 2 and 4). Several functional categories related to proteostasis were overrepresented in this synaptic type, including ‘Protein Stability’, or ‘Unfolded Protein Binding’, and Pura and Purg, involved in the transport of messenger RNA into the postsynapse^70^, were also found overexpressed. To further investigate this finding, we went back to the analysis of scRNAseq (Suppl. Table 3) and also found a very strong upregulation of most ribosomal genes in the Dentate Gyrus (Suppl. Fig. 8a). These findings, together with the recent discovery that local translation occurs at Mossy Fibre boutons^71^, indicate that proteostasis would play a particularly relevant role in this synaptic type.

### A unique extracellular matrix at EC-DG synapses

The proteome of EC-DG synapses presented several DE proteoglycans, including Bcan, Ncan, Agrn and Hspg2 (Vcan and Cspg5 also presented highest expression in EC-DG, but did not reach statistical significance, Fig. 3). The synaptic location of all these proteins is well documented^2^, mostly localizing to the extracellular matrix (ECM). Indeed, the GO term ‘Extracellular matrix structural constituent’ and the Reactome pathway ‘Integrin cell surface interactions’, related to the ECM, were overrepresented in EC-DG synapses. We thus observed a differential composition of the ECM, especially regarding the abundance of proteoglycans, that could specifically modulate the properties of this synaptic type. As previously, we also identified overexpressed proteins that are related to the regulation of AMPAR. These include the ‘receptor-type tyrosine-protein phosphatase delta’ (Ptprd)^72^, the AMPAR auxiliary protein Shisa9, first described in the DG^73^, and the scaffolding protein Epb41l1, known to bind to A1 subunits of the AMPAR^74,75^, regulating its activity-dependant insertion into the plasma membrane^76^.

Finally, proteins with highest expression in EC-DG synapses also retrieved several pathways related to the catabolism of branched chain amino, including ‘valine, leucine and isoleucine degradation’ (KEGG), ‘branched chain amino acid catabolism’ (Reactome) or ‘alpha amino acid metabolic process’ (GO). One of the two metabolic pathways to synthesize glutamate requires the catabolism of these amino acids, and the product of this reaction feed into the TCA cycle. EC-DG synapses might have a preferential use of this glutamate synthesis pathway, coupling synaptic transmission with energy production.

### Different neuronal expression of genes related to glutamate receptors

The fact that synapses formed by different neurons exhibit distinct expression patterns of proteins involved in the regulation of glutamate receptors prompted us to investigate whether this is mediated by genetic factors. Furthermore, as shown above, gene expression would explain some of the proteomic variability found between synapses (Fig. 2m-r). Thus, we returned to the ABCA^49^ database and explored gene expression in excitatory neurons of the hippocampus and subiculum. In these regions the ABCA defines 55 types of excitatory neurons, grouped into 8 classes. We first split all genes in two groups, those coding for our reference proteome, which we refer to as ‘synaptic genes’, and the rest (‘non-synaptic’ genes). We found that 18% of synaptic genes presented expression differences between neuronal classes (Suppl. Fig. 8b, Suppl. Fig. 9a, and Suppl. Table 5) and 17% between neuronal types (Suppl. Fig. 9b-i and Suppl. Table 6). Interestingly, the frequency of DE genes was 3 times higher among synaptic genes (Chi-square Test p < 0.0001, Suppl. Fig.8c). This remained significant if synaptic genes were compared to random gene sets of the same size taken from: i) all genes or ii) non-synaptic genes (Suppl. Fig. 8c).

Upregulated genes were mostly present in one neuronal class and eventually in two (Suppl. Fig. 8d), while downregulated ones appeared more repeatedly, in up to 5 classes (Suppl. Fig. 8e). The same happened in the comparison between neuronal types (Suppl. Fig. 8f), downregulated genes appeared more repeatedly. As our goal was to capture the functional categories most unique to each neuronal class/type, we only considered upregulated genes for subsequent analysis.

Next, we wanted to compare the expression patterns of upregulated synaptic genes between neuronal types. To achieve this, we computed expression correlation coefficients of these genes for each pair of neurons and performed hierarchical clustering. Neurons from the same class were grouped together (Fig. 4a), perfectly replicating the classification obtained by the ABCA with the entire transcriptome^49^. This indicates that synaptic genes from closely related neurons have more similar expression patterns, and that synaptic genes have a role in the classification of hippocampal neuronal types.

**Figure 4.**
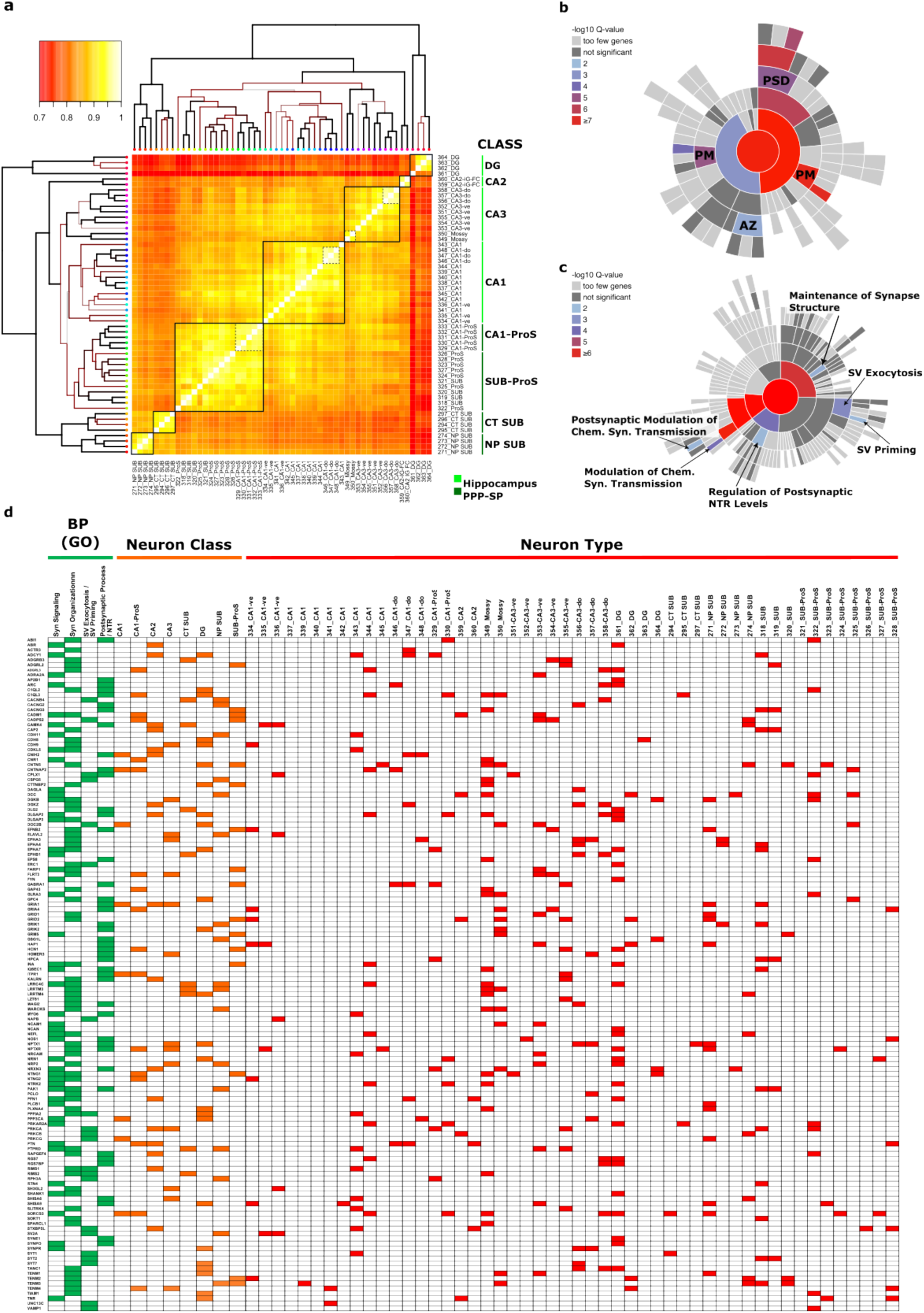
Differentially expressed genes mostly regulate neurotransmitter receptor function and synaptic vesicle exocytosis. **a.** Clustering of the coefficients of correlation for RNA expression of up-regulated genes with a synaptic location in excitatory neuron types from the hippocampal formation. **b.** Sunburst chart showing SynGO Cellular Component terms enriched among genes expressed at synapses that present increased expression in one or two types of excitatory neurons from the hippocampal formation. The background set for this analysis was the set of genes with a synaptic expression. Maximum stringency was applied for evidence filtering of SynGO annotations. PM: plasma membrane, AZ: active zone and PSD: postsynaptic density. **c.** Sunburst chart showing SynGO Biological Process terms enriched among genes expressed at synapses that present increased expression in one or two types of excitatory neurons from the hippocampal formation. **d.** Classes and types of excitatory neurons presenting increased expression of genes within Biological Process (GO) terms most overrepresented in the SynGO analysis.

To investigate common features among upregulated synaptic genes, we performed enrichment analysis of ‘Cellular Component’ and ‘Biological Process’ categories with the SynGO database. To obtain highly specific categories we used our reference proteome as a background set, and the most stringent criteria for evidence filtering. The first analysis found that these genes code for proteins residing in two main locations, the postsynaptic density (PSD) and the active zone (AZ) (Fig. 4b). The analysis of Biological Processes returned categories related to synaptic vesicle exocytosis and to the regulation of glutamatergic transmission, including the regulation of neurotransmitter receptor levels (Fig. 4c). Finally, we asked if the genes linked to these SynGO categories were spread across neuronal classes and types or if, instead, they were concentrated in a small number of them. We found that genes from these functional categories are widely spread across neuronal types (Fig. 4d), indicating that their differential regulation is a common trend among them.

We also investigated the signalling pathways associated with upregulated genes from individual neuronal types using pathfindR. In many instances the number of upregulated genes was small (Suppl. Table 6). Accordingly, pathfindR could only find enriched terms in 22 of the 55 neuronal types of the hippocampus and subiculum (Fig. 5a and Suppl. Table 7). Nevertheless, we observed that many of the enriched pathways were again related to the function of glutamate receptors (Fig. 5b). In 11 of the 22 types, upregulated genes were associated with pathways related to neurotransmitter receptor function, and in 8 this term was the most enriched one (Fig. 5b, dark blue bars). These included terms such as ‘ionotropic glutamate receptor activity’, ‘Trafficking of AMPA receptors’, ‘Activation of NMDAR and postsynaptic events’ or ‘Extracellular ligand gated ion channel activity’. In one neuronal type (CA1-343) the term ‘SV exocytosis’ was identified as the most enriched (Fig. 5b, red bars). These observations matched the findings obtained with SynGO (Fig. 4b,c), and strengthen them, as they were obtained with different databases and bioinformatic tools.

**Figure 5.**
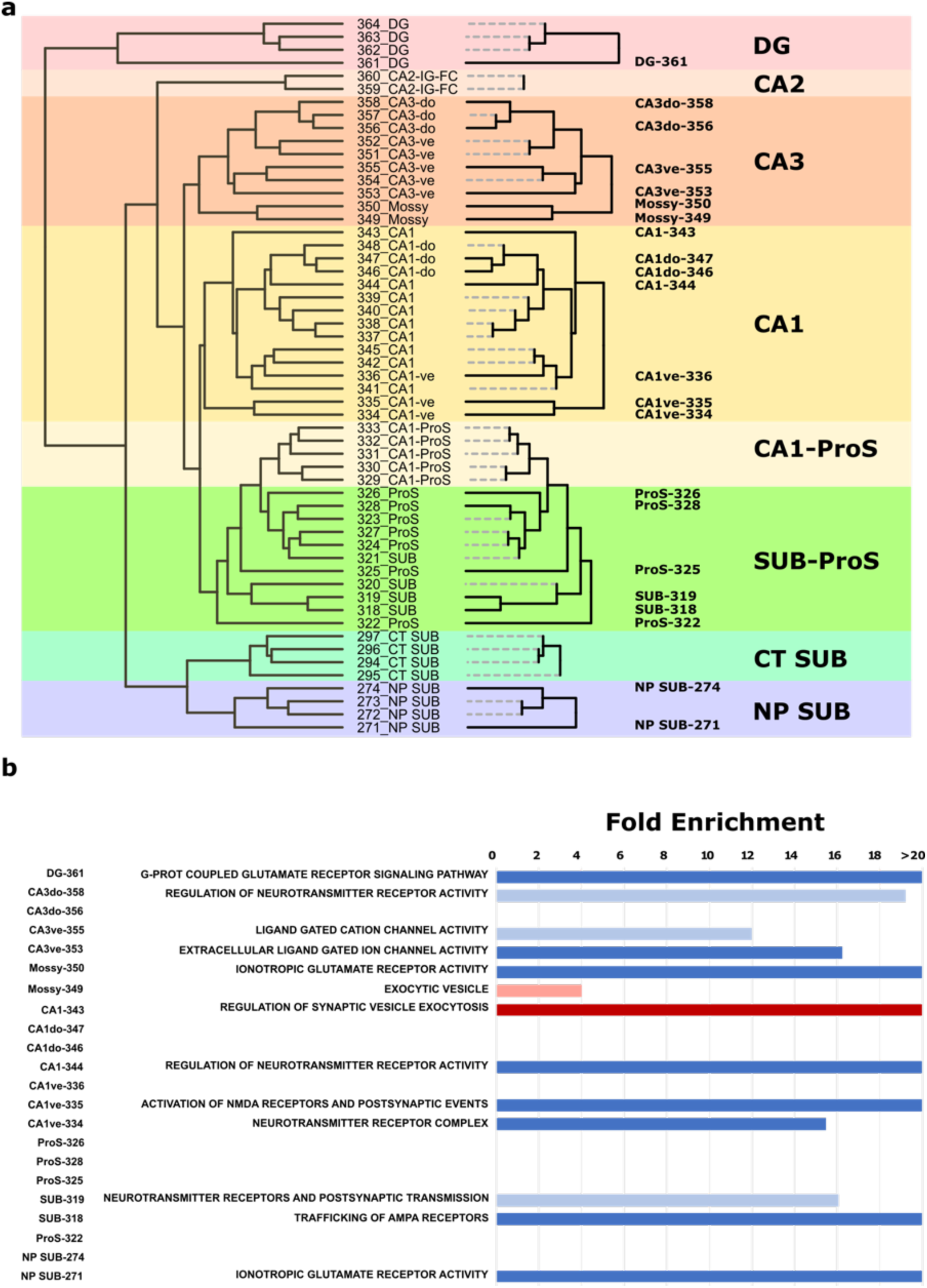
Hippocampal synaptic types are mostly defined by genes regulating neurotransmitter receptor function. **a.** Neuron types having genes expressed at synapses that show increased expression define neuron-specific synaptic types. Dashed lines correspond to neuron types whose upregulated genes cannot be linked to significantly overexpressed term. These synapses would not present any functional difference with those of other neurons from the same class. **b.** Fold enrichment of significantly enriched terms related to neurotransmitter receptor function (blue bars) or synaptic vesicle exocytosis (red bars). Dark blue or red denotes a term that is the most enriched one for that synaptic type. Light colours denote terms that are enriched but are not the most enriched. Fold enrichment corresponds to the ratio between the number of observed and expected genes related to one term.

### Genes related to glutamatergic signalling drive neuronal classifications

We found that synaptic genes present greater transcriptomic variation (Suppl. Fig. 8c), and that the ABCA neuronal classification^49^ can be replicated considering only synaptic genes (Fig. 4a). To investigate if synaptic genes contribute to the classification of hippocampal neurons, we referred again to the ABCA database, and first confirmed we could replicate their classification, as indicated by the segregation of neuronal classes in nonlinear dimensionality reduction maps (U-Map) (Fig. 6a). Noticeably, the U-map made with synaptic genes (Fig. 6b) was highly similar to that produced with all genes. Instead, U-Maps from non-synaptic genes had very different topologies, with high overlaps between neurons from different classes (Fig. 6c and Suppl. Fig. 10a). This suggests that synaptic genes drive the classification of hippocampal excitatory neurons, as it has been shown for cortical neurons^77^. To validate this observation, we used the Random Forest method, a supervised machine learning approach^78^, that computes the relative contribution of each gene to the classification.

**Figure 6.**
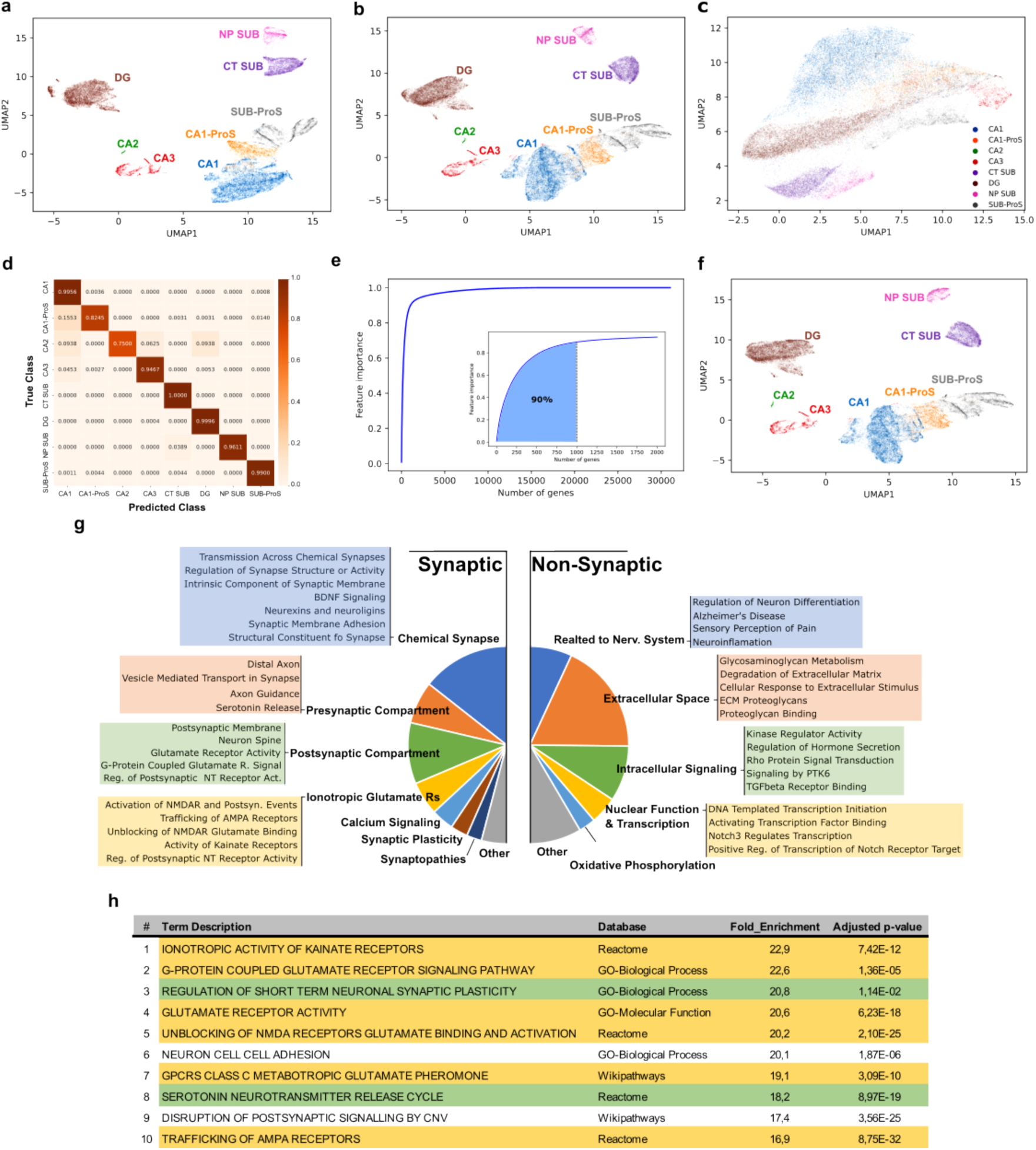
Expression differences in genes encoding synaptic proteins strongly determine the classification of excitatory neurons. **a.** UMAP graph generated with single-cell RNA abundance data obtained from excitatory neurons in the 8 classes identified in the hippocampal formation. Abundance of all genes in the genome was considered for the construction of this graph. ProS, prosubiculum; SUB, subiculum, NP SUB; near-projecting neurons from the subiculum and CT SUB; corticothalamic neurons from the subiculum. **b.** UMAP generated as in (a), although in this occasion only genes coding for synaptic proteins were considered. **c.** UMAP generated as in (a), using a random set of genes not expressed at synapses, with the same number of genes as in the synaptic dataset in (b). **d.** Confusion or error matrix generated by the Random Forest algorithm, showing the success rates in assigning a class to each neuron. Colour legend correspond with the accuracy of the prediction, 1 being perfect accuracy. **e.** Cumulative importance of the expression level of each gene in the genome for the classification of excitatory neurons into classes. Inset, cumulative Importance of the top 2000 genes with the highest importance to the classification. Note that the top 1000 contributing genes provide 90% of the information necessary to construct the classification. **f.** UMAP generated as in (a) but using only the 520 synaptic genes found among the Top 1000 genes contributing to the classification. **g.** Main signalling pathways and biological functions found among genes encoding for synaptic and non-synaptic proteins of the top 1000 that most contribute to the classification of excitatory neurons into classes. **h.** Top 10 signalling pathways with the largest fold enrichment. In yellow those relative to the function of ionotropic or metabotropic glutamate receptors. In green those relevant to presynaptic function.

After the training phase, the algorithm could predict neuronal classes with high accuracy (total accuracy for the train set 0.9893 - total accuracy for the test set 0.9014), indicating that the computed contribution of each gene to the classification was reliable. Indeed, the predictive power of the algorithm was above 95% for 6 of the 8 neuronal classes (Fig. 6d). We found that a small number of genes did drive the overall classification. The added weight of the top 1000 genes contributing to the classification accounted for 90% of the information carried by the whole transcriptome (Fig. 6e and Suppl. Table 8). Importantly, over 50% of these genes were synaptic (Suppl. Fig. 10b), a 4x overrepresentation that was highly statistically significant (Chi-square test, p < 1e-23). Using the synaptic genes in the top 1000 was sufficient to replicate the U-map generated with the entire transcriptome (Fig. 6f). Furthermore, the accuracy of the Random Forest prediction was better when using synaptic genes as opposed to the entire transcriptome and best when using the synaptic genes found in the top 1000 list (Suppl. Fig. 10c).

Using the Chi-square stat, we found that genes expressed at synapses were more over-represented in genes driving the classification than genes enriched in the PSD^79^, in the MASC complex^80^ or in other functional categories enriched in the top 1000 genes contributing to the classification (Suppl. Fig. 10d). Random Forest performance was also good in classifying neurons into types, although less accurately (total accuracy of the train set 0.8559 and total accuracy of the test set 0.7653, Suppl. Fig. 10e). The list of the top 1000 genes most relevant to the classification of types also carried over 90% of the weight, and included over 500 synaptic genes (Chi-square test, p < 1e-10, Suppl. Fig. 10b and Suppl. Table 8). pathfindR analysis of synaptic genes in the top 1000 contributing to the classification of neuronal classes revealed synaptic functions or locations related to both pre and postsynaptic compartments (Fig. 6g-h and Suppl. Table 8). Yet, terms with highest fold enrichments were mostly related to the function and organization of glutamate receptors (Fig. 6h). Curiously, non-synaptic genes of the top 1000 were also associated with some functions of the nervous system (i.e. Neuron differentiation or Neuroinflammation), among others.

## Discussion

Electrophysiological studies have shown that different synaptic types have unique functional properties^81,82^. Yet the molecular basis driving these differences are poorly understood. Investigating synaptic types at the proteomic level has been challenging, as they are confined to microscopic brain regions. To address this limitation, we have developed a procedure to obtain microscopic brain samples, and to extract synaptic proteins from them in sufficient quantities for high-throughput proteomics. This method has several advantages, it provides a great level of anatomical resolution, as the location of the collected samples is known; it delivers a wide and deep coverage of the synaptic proteome, identifying proteins from most subsynaptic compartments and it can be used in any species, including humans, as it does not require prior genomic manipulations. We have used this approach to profile the proteome of the synapses constituting the trisynaptic circuit of the hippocampus. Its anatomical organization segregates these synapses into different layers where they account for close to 90% of all^29–38^.

A relevant conclusion of our proteomics study is that essentially the same proteins are present in all synapses investigated. This observation denotes that functional diversity arises from changes in the abundance of shared components. These changes would result in specific molecular processes being favoured at individual synapses. For example, it is well-known that CA3-CA1 synapses require the activation of NMDARs for LTP expression but DG-CA3 synapses don’t. Several synaptic types express forms of NMDAR-independent LTP across the brain, and class I metabotropic glutamate receptors (Grm) are involved in some of them^83,84^. Indeed, the role of Grm1/5 in NMDAR-independent LTP at DG-CA3 synapses has been addressed by a few studies, albeit these returned contradictory results^17^. Our data provides strong support for a role of Grm1 in LTP at DG-CA3 synapses, as this receptor and several of its downstream signalling molecules are highly expressed in them. Thus, while these molecules are present in both synaptic types, their increased abundance in DG-CA3 synapses could allow them to express an NMDAR-independent LTP, finetuning the functional properties of this synaptic type.

Differentially expressed proteins were involved in many signalling pathways and biological processes relevant to synaptic biology. Remarkably, most of them were exclusively found in one synaptic type, suggesting they could contribute specific functions. CA3-CA1 synapses exhibited several overrepresented pathways directly related with AMPAR trafficking, particularly regarding GluA2-containing AMPARs, and clathrin mediated endocytosis, the primary mechanism by which AMPARs are removed from synapses^58^. Proteins in CA3-CA1 synapses also displayed many functional categories related to actin polymerization and branching, key processes in spine structural plasticity. The non-canonical Wnt/Ca^2+^ pathway, which regulates calcium release from internal stores^61^, was also overrepresented in this synaptic type. And so were numerous metabolic pathways related to energy production, suggesting they might have increased energetic demands.

Instead, DG-CA3 synapses were characterised by signalling pathways downstream of class I metabotropic glutamate receptors. They also exhibited a striking increase in ribosomal proteins, likely due to an elevated number of presynaptic ribosomes, as protein translation at mossy fibre boutons would regulate synaptic plasticity^71^. They also presented increased levels of proteins that positively regulate ERK1/2 signalling, a pathway linking ionotropic glutamate receptors with protein translation. In line with previous findings, showing that mossy fibre boutons have the highest level of ERK1/2 activation in the hippocampus^85^. Furthermore, DG-CA3 synapses presented increased abundance of all 4 proteins organizing intermediate neurofilaments. These proteins have been confidently identified in synapses^68^, being involved in synaptic transmission and plasticity^68^. Our data indicates that CA3-CA1 and DG-CA3 synapses would have specific requirements regarding their cytoskeletal requirements. Structural plasticity studies at dendritic spines show considerable differences between neurons, which might originate in different cytoskeletal compositions^86^. Finally, EC-DG synapses were characterised by a unique ECM, with increased levels of several proteoglycans and other of its constituents. The synaptic localization of proteoglycans is also well documented^2^, contributing to AMPAR trafficking^87,88^ and synaptic transmission^89^. Indeed, the ECM is known to restrict AMPAR mobility^90^.

Overall, our proteomic findings provide support for considerable molecular diversity among the synapses of the trisynaptic loop. Impacting multiple domains of synaptic biology, including the trafficking and synaptic stability of AMPARs, spine structural plasticity, signalling through metabotropic receptors, control of calcium levels, local protein translation or regulation of the energetic metabolism, among others. Importantly, however, we also identified DE proteins controlling the function of glutamate receptors in all samples studied. As we had seen that gene expression contributes to synaptic proteome diversity, we explored if gene expression mechanisms could be involved in this common feature.

We found that synaptic genes differentially expressed between neuronal types mostly localized to two subsynaptic locations, the active zone, and the postsynaptic density. Being involved in synaptic vesicle (SV) exocytosis, and the postsynaptic regulation of chemical synaptic transmission, especially in the regulation of neurotransmitter receptor levels at the synapse. Importantly, genes involved in these processes had differential expression patterns in most neuronal types from the ABCA, with each type overexpressing a subset of them. Therefore, the differential expression of these genes would be a common trend among excitatory neurons in the hippocampus and subiculum. In a second analysis we investigated the signalling pathways overrepresented in independent neuronal types. These analyses also retrieved many pathways related to glutamate receptor function, these being the most enriched ones for many neuronal types. While pathways related to SV exocytosis were weakly overrepresented in this analysis. An orthogonal mathematical approach based on machine learning corroborated the differential expression of genes related to glutamatergic function between neuronal types. This approach was employed to identify the genes that contribute the most to transcriptomics-based neuronal classifications. Showing that genes involved in glutamatergic function were key to these classifications.

In the present study, we introduce a procedure that allows to explore the synaptic proteome of anatomically defined microscopic brain regions. With this method we have been able to identify major molecular differences between the synapses that comprise the trisynaptic circuit. This is an important resource to advance in our understanding of the molecular mechanisms controlling their diverse functional properties. More importantly, our combined investigation of proteomic and transcriptomic datasets indicates that glutamate receptors and proteins directly controlling their function, are common drivers of synaptic proteome variability, possibly having key contributions to their specific properties. Remarkably, neuron-specific transcriptional mechanisms would contribute to the unique expression levels of these synaptic proteins.

## Methods

### Animal handling

All animal research was done with C56BL/6J mice (Jackson Laboratories, Research Resource Identifier, RRID:MGI:5656552) in accordance with national and European legislation (Decret 214/1997 and RD 53/2013). Research procedures were approved by the Ethics Committee on Animal Research from the: i) Institut de Recerca de ĺHospital de la Santa Creu i Sant Pau (IR-HSCP) and from the ii) Universitat de Barcelona for whole-cell recording experiments. These procedures were also approved by the Departament de Territori i Sostenibilitat from the Generalitat de Catalunya (approval reference numbers 9,655 and 164.16). Maintenance and experimental procedures were conducted at the animal facilities of the IR-HSCP or the Faculty of Medicine of the Universitat de Barcelona, for whole-cell recording experiments. Mice were housed at a 12h light/dark cycle, with fresh water and food ad libitum. We used animals of both sexes and 9-14 weeks of age. 12 animals were used for laser-capture microdissection proteomics experiments, 2 to isolate postsynaptic density fractions using sucrose gradients and 12 for manual hippocampal dissection and preparation of triton insoluble membranes. 12 animals were used for double immunofluorescence in brain sections. Finally, 22 mice were used for electrophysiological studies.

### Mouse brain dissection

Mice were culled by cervical dislocation, the head was dissected, and brain removed from skull and meninges. All brain dissection manipulations were done in the presence of chilled 1x phosphate-buffered saline (PBS, 0.144 M NaCl, 2.683 mM KCl, 10.144 mM Na_2_HPO_4_, 0.735 mM KH_2_PO_4_, [P5368-10PAK from Sigma]). Cerebellum and olfactory bulb were removed prior to any other manipulation. For laser-capture microdissection the forebrain was wrapped in aluminium foil, snap frozen in liquid nitrogen and stored at −80C. For isolation of postsynaptic density (PSD) fractions by ultracentrifugation hippocampi were dissected using iris scissors (PMD120; Thermo Scientific), tissue forceps 1:2 (PMD023445; Thermo Scientific) and scalpel blades in chilled glass petri dishes. Entire hippocampi were frozen at −80C before processing. For manual dissection of CA1, CA3 and DG regions readily dissected hippocampi were first cut coronally in 500 μm slices in the presence of chilled 1x PBS using a tissue slicer (Kerr Scientific Instruments). 8-12 slices where obtained from each hippocampus. Slices were immediately transferred into a glass petri-dish with chilled 1x PBS using a small paint brush. Next CA1, CA3 and DG regions were manually separated from each other using 18G needles (BD) under a microscope Carl Zeiss Meditec model S100/OPMI 1-FC (see Supplementary Video for a demonstration of manual dissection of hippocampal regions). Dissected regions were placed in individual tubes containing chilled homogenization buffer with phosphatase and protease inhibitors (0,32M Sucrose; 10mM HEPES pH 7,4; 2mM EDTA; 5mM sodium o-vanadate; 30mM NaF; 2μg/ml aprotinin; 2μg/ml leupeptin and 1:2000 PMSF (v/v)) with a pasteur pipette and frozen dry at −80C.

### Laser-capture microdissection of hippocampal neuropile

Frozen forebrains were used to obtain 10 μm thick coronal sections in a Leica CM1950 cryostat. Only sections that contained the dorsal hippocampus (Suppl. Fig. 1a) were processed by laser-capture microdissection. Sections were placed in membraneSlide 1.0 PEN microscope slides (Zeiss, 415190-9041-000) and stored at −20C. The neuropile of CA1, CA3 and dorsal DG were microdissected using a Leica LMD 6000 laser microdissection microscope. Between 90 and 110 mm^2^ were microdissected for each hippocampal region and biological replica. Three biological replicas were generated for each area. All microdissected tissue for each replica was collected in the same 1.5ml tube.

### Isolation of synaptic fractions from laser-capture microdissected tissue

Laser-capture microdissected tissue was collected in 1.5 ml tubes and mixed with PBS containing 1% Triton X-100, 2μg/ml leupeptin and 1/2500 PMSF. The sample was then sonicated in an ultrasonic bath (Branson 1510) for 2 min, incubated in agitation (300rpm) in a ThermoMixer C (Eppendorf) for 30 min at 35C and sonicated again as previously. Afterwards, sample was centrifuged for 10 min at 21.000xg at 4C in a refrigerated centrifuge (Eppendorf 5417R). The pellet was resuspended in PBS with 1% SDS. The resuspended pellet and supernatant were mixed with 10x SDS sample buffer for analysis by proteomics or immunoblot. Tissue extraction was also performed with a RIPA buffer containing PBS, 0.1% SDS, 0.5% sodium deoxycholate and 2% Triton X-100 buffer.

### Isolation of synaptic fractions from hippocampal subfields

Manually dissected hippocampal subfields (CA1, CA3, dDG; see Supplementary Video) from 3 animals where accumulated for each biological replica. A total of four biological replicas were prepared for each region. CA1 samples were homogenized in 450μl of homogenizing buffer (HB), CA3 and DG in 300μl. Homogenizing buffer composition: 0,32M Sucrose; 10mM HEPES pH 7,4; 2mM EDTA; 5mM sodium o-vanadate; 30mM NaF; 2μg/ml aprotinin; 2μg/ml leupeptin and 1:2000 PMSF (v/v). Homogenization was performed in 1ml borosilicate tissue homogenizers (357538, Wheaton), using 20-30 strokes. The homogenate was centrifugated in 1.5ml tubes at 800xg and 4C for 10 min in a Eppendorf refrigerated centrifuge (Eppendorf 5417R). The pellet, containing the nuclear fraction and cell debris, was re-homogenized once in the same buffer and centrifuged in the same conditions. Supernatants from both centrifugations were pooled and spun down at 10.000xg for 15 min at 4C in the same centrifuge. The resulting pellet was resuspended in Triton buffer (TB: 50mM HEPES pH7.4; 2mM EDTA; 5mM EGTA; 1mM sodium o-vanadate; 30mM NaF; 1% Triton X-100; 2μg/ml aprotinin; 2μg/ml leupeptin and 1:2000 PMSF (v/v)). TB volume used was ½ HB. This mixture was left in ice for 15 minutes and centrifuged at 21.000xg for 30 min at 4C in the same centrifuge. The resulting pellet was resuspended with 30μl of 50mM Tris pH 7.1; 1% SDS and incubated with this buffer for 15 min at room temperature. A final centrifugation was done at 21.000xg for 15 min at room temperature. The resulting supernatant corresponds with the postsynaptic density enriched fraction.

### Isolation of synaptic fractions by differential ultracentrifugation

Isolation of synaptic fractions using differential ultracentrifugation involves first the separation of synaptosomes on the bases of their sedimentation rate in sucrose density gradients and the isolation of synaptic protein complexes insoluble to the non-ionic detergent Triton X-100^6,9,79^. Briefly, the hippocampi from two mice were homogenized in 1ml borosilicate tissue homogenizer (357538, Wheaton) adding 9ml of homogenizing buffer for each 1g of tissue weight. Homogenization was done with 20-30 strokes. Homogenizing buffer composed of: 0,32M Sucrose; 10mM HEPES pH 7,4; 2mM EDTA; 5mM sodium o-vanadate; 30mM NaF; 2μg/ml aprotinin; 2μg/ml leupeptin and 1:2000 PMSF (v/v). This sample was first centrifuged at 1400xg and 4C for 10 minutes in a refrigerated centrifuge (Eppendorf 5417R). The pellet of this centrifugation was re-homogenized twice following the same procedure. The three supernatants generated were pooled and centrifuged at 700xg for 10 minutes, the pellet was discarded. Next, the sample was centrifuged at 21.000xg for 30 minutes at 4C in the same centrifuge. The resulting pellet was resuspended with Tris 50mM pH7.4 and 0,32M sucrose. A sucrose gradient was prepared with 1 ml of (top to bottom): sample; 50 mM Tris pH 7.4, 0.85 M sucrose; 50 mM Tris pH 7.4, 1 M sucrose; 50 mM Tris pH 7.4, 1.2 M sucrose. This gradient was centrifuged in a SW60Ti rotor (Beckman Coulter) at 82.500xg for 2 hours. The 1.0-1.2 interphase was collected, diluted with 2 equal volumes of 50mM Tris pH 7.4, and centrifuged at 21.000xg for 30 minutes at 4C. The subsequent pellet was resuspended in 50mM Tris pH 7.4, 1% Triton X-100 and maintained in ice for 10 min. This sample was centrifuged at 21,000xg during 30 min at 4C, the resulting pellet corresponds with the final synaptic fraction.

### Protein electrophoresis and immunoblot

Sample preparation for protein electrophoresis and immunoblot was accomplished by mixing it with 10x SDS loading sample buffer, composition: 500mM Tris pH7.4; 20% SDS; 50% glycerol and 10% b-mercaptoethanol. Prior to its analysis samples were boiled at 95C for 5 min.

SDS-PAGE gels were run in a vertical MiniProtean system kit (Bio-Rad) with 1x running buffer (25 mM TRIS pH 8.4; 0.187 M glycine and 0.1% SDS). Protein standards used were All blue Precision Plus (Bio-Rad). For LC-MS/MS analysis protein gels were stained over night at room temperature with Coomassie solution (B8522-1EA; Sigma-Aldrich) and washed with 2.5% acetic acid and 20% methanol and subsequent washes of 20% methanol, until protein bands were clearly visible. For immunoblot TGX Stain-Free™ gels (161-0181 & 161-0185, SF gels; Bio-Rad) were used and activated as recommended by the manufacturer. Gel images were acquired with ChemiDoc XRS+ (Bio-Rad) and quantified with Image Studio Lite ver. 3.1 (LI-COR Biosciences).

Protein transference was done using a MiniProtean kit (Bio-Rad), and 1x chilled transference buffer (20% methanol; 39 mM Glycine; 48 mM TRIS; 0.04% SDS). Proteins were transferred onto methanol pre-activated polyvinylidene fluoride (PVDF) membranes (IPFL00010, Immobilon-P; Merck-Millipore). Membranes transferred from TGX Stain-Free™ gels were imaged and quantified for posterior normalization with a ChemiDoc XRS+ (Bio-Rad) using the Image Lab software (Bio-Rad). After transference, PVDF membranes were blocked with 5ml Odissey blocking solution (927-50000; LI-COR) diluted with 1x tris-buffered saline (TBS, 50 mM Tris pH7.4; NaCl 150mM and 0.1% sodium azide). Next, membranes were incubated with primary antibodies in Tween-TBS (T-TBS: 0,1% Tween 20 - TBS) ON at 4C or 1 hour at room temperature. Primary antibodies used: Psd95 (#3450; Cell Signaling, [RRID:AB_2292883]); Synaptophysin (Ab8049; Abcam [SY38], [RRID:AB_2198854]); GluA2 (MAB397; Millipore [RRID:AB_2113875]; Shisa6 (NBP2-85726; Novus Biologicals [RRID:AB_3427376]); mGluR2 (# 191 103; Synaptic Systems [RRID:AB_2232859]; Prkar2a (ab32514; Abcam [RRID:AB_777289]); Ptprd (NBP2-94767; Novus Biologicals [RRID:AB_3464681]). Antibody dilution was 1:1000 except for mGluR2, Ptprd, Prkar2a (1:500) and Shisa6 (1:250). Membranes were washed four times with 1x T-TBS for 5 min before incubation for 1 hr at room temperature protected from light with 5 ml of the following secondary antibodies prepared in T-TBS at a dilution of 1:7.500: anti-rabbit (926-68073, IRDye 680CW, [AB_10954442]), anti-mouse (926-32212, IRDye 800CW [RRID:AB_621847] or 925-68072, IRDye 680RD, [RRID:AB_2814912]) and anti-goat (926-32214, IRDye 800CW, [RRID:AB_621846]). Images were acquired with an Odissey Scanner (LI-COR Biosciences) and protein bands were analysed with Image Studio Lite ver. 3.1 (LI-COR Biosciences). Protein abundance in postsynaptic density enriched fractions was normalized by the abundance of Psd95, a marker of postsynaptic densities, to correct for purity differences between samples.

### Sample processing for mass spectrometry

Synaptic fractions obtained from laser-captured microdissected tissue or PSD fractions generated with standard procedures were analysed by conventional protein gel electrophoresis in 6% polyacrylamide gels. For LCM samples gels were run to half their length and stained with Coomassie as described above. After distaining LCM samples were cut into 5 bands of the same size (Suppl. Fig. 2h). PSD samples were separated into 13 electrophoretic bands (Suppl. Fig. 2i). Next, gel bands were cut into 1×1 mm cubes with a scalpel blade in an ethanol cleaned glass plate and under a laminar flow hood. Gel cubes were transferred to 1.5ml tubes for proteomic analysis (0030 123 328; Eppendorf). 50 mM ammonium bicarbonate (ABC) in 50% ethanol was added to each tube and incubated for 20 min at room temperature. This solution was replaced with absolute ethanol and incubated 15 more min. For protein reduction gel cubes were mixed with freshly prepared 10mM DTT (dithiothreitol; Merck) in 50mM BA and incubated 1 h at 56C. For protein alkylation, DTT was removed and freshly prepared 55mM IAA (iodacetamide; Merck) in 50mM BA added, incubation was performed in the dark for 30 min at room temperature. IAA was removed, 25mM BA added to gel cubes and incubated in the dark for 15 min. For in-gel protein digestion reduced and alkylated samples were mixed with 25 mM BA-50% acetonitrile (ACN) and incubated 15 min twice. Gel cubes were dehydrated with 100% ACN for 10 min. Next, trypsin (Promega) containing solution was prepared and incubated with gel cubes ON at 30C. Tryptic peptides were extracted from gel cubes by first adding 100% ACN and incubating 15 min at 37C. Later, 0.2% trifluoroacetic acid (TFA) was added and incubated for 30 min. Supernatants were transferred to 0.5 ml tubes (#0030 123 301; Eppendorf) previously washed with ACN to prevent peptide binding to the walls. Liquid-phase was evaporated using a SpeedVac (Thermo-Fisher Scientific). Dried peptides were resuspended in 5% ACN and 0.1% formic acid and bath sonicated for 2 min. Samples were then centrifuged at maximum speed to remove possible gel remainings. Samples were stored at −20C.

### Mass spectrometry analysis of tryptic peptides

Tryptic peptides were analysed by LC-MS/MS using an EASY-nLC system (Proxeon Biosystems, Thermo Fisher Scientific) connected to a Velos-Orbitrap mass spectrometer (Thermo Fisher Scientific, Bremen, Germany). Instrument control was performed using Xcalibur software package, version 2.1.0 (Thermo Fisher Scientific, Bremen, Germany). First, peptide mixtures were fractionated by on-line nanoflow liquid chromatography with a two-linear-column system. Digests were loaded onto a trapping guard column (EASY-column, 2 cm long, ID 100 μm, packed with Reprosil C18, 5 μm particle size from Proxeon, Thermo Fisher Scientific) at a maximum pressure of 160 Bar. Then, samples were separated on the analytical column (EASY-column, 10 cm long, ID 75 μm, packed with Reprosil, 3 μm particle size from Proxeon, Thermo Fisher Scientific). Elution was achieved by using a mobile phase from 0.1% formic acid and 100% acetonitrile with 0.1% formic acid and applying a linear gradient from 5 to 35% of buffer B for 120 min at a flow rate of 300 nL/min. Ions were generated applying a voltage of 1.9 kV to a stainless-steel nano-bore emitter (Proxeon, Thermo Fisher Scientific), connected to the end of the analytical column. The LTQ Orbitrap Velos mass spectrometer was operated in data-dependent mode. A scan cycle was initiated with a full-scan MS spectrum (from mass to charge [m/z] 300 to 1600) acquired in the Orbitrap with a resolution of 30,000. The 20 most abundant ions were selected for collision-induced dissociation fragmentation in the linear ion trap when their intensity exceeded a minimum threshold of 1000 counts, excluding singly charged ions. Accumulation of ions for both MS and MS/MS scans was performed in the linear ion trap, and the AGC target values were set to 1 × 10^6^ ions for survey MS and 5000 ions for MS/MS experiments. The maximum ion accumulation time was 500 and 200 ms in the MS and MS/MS modes, respectively. The normalized collision energy was set to 35%, and one microscan was acquired per spectrum. Ions subjected to MS/MS with a relative mass window of 10 ppm were excluded from further sequencing for 20 s. For all precursor masses a window of 20 ppm and isolation width of 2 Da was defined. Orbitrap measurements were performed enabling the lock mass option (m/z 445.120024) for survey scans to improve mass accuracy.

LC-MS/MS data was analysed using Progenesis QI software (Nonlinear Dynamics, Newcastle, UK). This software allows to review the chromatogram alignments, to filter the data, to review peak picking, to normalize the data and to identify peptides, among other features. Specifically, sample ions were automatically aligned to compensate for drifts in retention time between runs. Yet, they were also reviewed and edited manually. The peak picking limits were automatic, the main ion charge selected was set at 4 and the retention time limits were adjusted according to the chromatograms in each sample. Peptide ions were filtered by removing those with a charge of 1 or >4, *m/z* from 300 to 1,600 and the specific retention determined for each case was also set. Progenesis was also used to normalize peptide and protein abundances, allowing for sample comparisons. Log of abundance ratios between each LC-MS/MS run and a reference run from the same dataset, which is selected by the Progenesis algorithm, are first computed. Next, the median of the log ratios is calculated for each of the runs. The variance of the ratio distribution is also used iteratively to remove outliers. Finally, the ratio between the median calculated for each run and the reference run is used as a scalar factor for recalibration of all runs.

### Database search of mass spectrometry data

All MS/MS samples were analysed using Mascot (Matrix Science, London, UK; version“2.5.1). Mascot was searched with a fragment ion mass tolerance of 0.80 Da and a parent ion tolerance of 10.0 ppm. Charge state deconvolution and deisotoping were not performed. MS/MS spectra were searched with a precursor mass tolerance of 10 ppm, fragment tolerance of 0.5-0.8 Da, trypsin specificity with a maximum of 2 missed cleavages, cysteine carbamidomethylation set as fixed modification (up to 57) and methionine oxidation as variable modification (up to 16). The quantification method applied to quantify protein abundances was a label-free based approach.

### Criteria for protein identification by mass spectrometry data

Scaffold (version Scaffold_4.8.5, Proteome Software Inc., Portland, OR) was used to validate MS/MS based peptide and protein identifications obtained from Mascot. Peptide identifications were accepted if they could be established at greater than 95,0% probability by the Peptide Prophet algorithm^91^ with Scaffold delta-mass correction. Protein identifications were accepted if they could be established at greater than 99,0% probability and contained at least 2 identified peptides. Protein probabilities were assigned by the Protein Prophet algorithm^92^. Using these filters a protein false discovery rate (FDR) under 1.0 was achieved, at the level of the entire dataset, as estimated by a search against a target-decoy database. Proteins that contained similar peptides and could not be differentiated based on MS/MS analysis alone were grouped to satisfy the principles of parsimony.

### Peptide and protein quantification

Peptide abundances were calculated and normalized using Progenesis, which integrates the area under the curve (AUC) of MS1 peaks for peptide quantification. Normalized peptide abundances were exported from Progenesis and peptides from proteins not identified by Scaffold were discarded. Next unique peptides were identified as those defined as non-conflicting by Progenesis or identified as unique by NextProt tool (Expasy) or the Peptide Search tool from Uniprot. Abundances from species of the same unique peptide identified with different retention times were added together. Abundances from modified peptides were added separately. Finally, peptide abundances were normalized based on the average abundance of all peptides from the 14 main postsynaptic density (PSD) scaffolds (Dlg1, Dlg2, Dlg3, Dlg4, Dlgap1, Dlgap2, Dlgap3, Dlgap4, Shank1, Shank2, Shank3, Homer1, Homer2 and Homer3), thus correcting for synaptic enrichment differences between purifications. Peptide abundances were then analysed with MSqROB to obtain protein abundance data and to identify proteins differentially expressed between groups^41,42^. MSqROB was used with the following settings: abundance data was log2 transformed, no normalization was applied, each peptide had to be identified in at least two experiments and only proteins identified with at least 2 peptides were considered for quantification. Furthermore, genotype was used as the fixed effect, while run, sequence and peptide modification were defined as random effects.

### Tissue processing for double immunofluorescence analysis

Mice were first anesthetized with a solution containing Ketamine (120 mg/kg) and Xylazine (30 mg/kg). Next, an intracardial perfusion of 4% paraformaldehyde solution in 0.1 M phosphate buffer (pH 7.4) was performed. Brains were extracted, post-fixed overnight in the same solution and then cryoprotected in a 30% sucrose solution in 0.1 M phosphate buffer for 48 h at 4°C. Subsequently, brains were frozen in ice-cold 2-methylbutane (Merck, 1060561000) and stored at −80°C. Coronal brain sections, ranging from bregma −1.46 mm to bregma −2.06 mm, were prepared using a Leica CM1950 cryostat. Free-floating sections, 25 μm thick, were preserved at −20°C in a 0.01 M antifreeze solution containing 20% sucrose, 30% ethylene glycol, and 1% polyvinylpyrrolidone (PVP) until used.

### Double immunofluorescence in adult mouse brain sections

Frozen free-floating brain sections were first washed with PBS (0.1 M, pH 7.4) and next with PBS containing 0.1% Triton X-100 to remove the antifreeze solution. The sections were then incubated at room temperature (RT) for 1 h in a blocking buffer containing 10% foetal bovine serum (FBS), 3% bovine serum albumin (BSA) and 0.25% Triton X-100 in PBS. After blocking, sections were incubated overnight at 4°C with the primary antibody corresponding to the protein of interest diluted in blocking buffer. Following 0.1% Triton X-100 PBS washes, sections were incubated for 1 h at 37°C with the corresponding secondary antibody diluted in blocking buffer. Subsequently, sections were incubated for 2 h at 37°C with the primary antibody for the pre-(vGlut1) or post-synaptic (Psd95) marker, followed by additional washes and a final incubation with its corresponding secondary antibody. Following this procedure nuclei were stained with 4’,6-diamidino-2-phenylindole (DAPI) (1:10,000; D9542, Sigma-Aldrich) in PBS during 10 min at RT. Sections were washed with PBS, mounted on slides, and coverslipped using ProLong Glass anti-fade mounting medium (P36984, Thermo Fisher Scientific). Representative images of the hippocampus were captured using a Leica inverted fluorescence confocal microscope (Leica TCS SP5-AOBS, Wetzlar, Germany) with an HCX PL APO 63x oil/0.6-1.4 objective. To minimize crosstalk and bleed-through effects, sequential scanning was employed. Fluorescent images were acquired in a 1024×1024 pixel scan format within a spatial dataset (xyz) and processed using Leica Standard Software TCS-AOBS. Confocal images were analysed with the software FIJI/ImageJ^93^ to quantify the signal from the pre- or post-synaptic marker overlapping with the protein of interest. Using the ‘Image Calculator’ function we generated an image that represents the overlap from both channels and measure the Integrated Density (IntDen) of the overlapping signal. These values were then compared between images from the three different hippocampal layers.

Primary antibodies used: Psd95 (1:100 dilution, Thermofisher MA1045; [RRID:AB_325399]), vGlut1 (1:750, Merck Millipore AB5905; [RRID:AB_2301751]), Homer2 (1:100, 160203, Synaptic Systems; [RRID:AB_10807099]), Calcineurin (1:100, 387002, Synaptic Systems; [RRID:AB_2661875]), Homer3 (1:100, 160303, Synaptic Systems; [RRID:AB_10804288]), Synaptoporin (1:100, 102003, Synaptic Systems; [RRID:AB_2619748]), Epb41l1 (1:100, 276103, Synaptic Systems; [RRID:AB_2620007]) and Mpp2 (1:50, HPA073483, Merck, [RRID: AB_3678682]).

Secondary antibodies used: Anti-Rabbit Alexa-Fluor-488 (1:500, 160203, Invitrogen), anti-Mouse Alexa-Fluor-594 (1:500, 387002, Invitrogen) and anti-Guinea pig Alexa-Fluor-647 (1:1000, AB5905, Invitrogen).

### Slice preparation for whole-cell electrophysiological recordings

Hippocampal acute slices were prepared from 8 to 12 weeks old C57BL/6J mice as follows. Animals were anesthetized with isoflurane and decapitated prior quick brain removal. Brains were then immersed in ice-cold artificial cerebral spinal fluid 1 (aCSF1 containing in mM, 206 sucrose, 1.25 NaH_2_PO_4_, 26 NaHCO_3_, 1.3 KCl, 1 CaCl_2_, 10 MgSO_4_, 11 glucose, purged with 95 % O2/5% CO2, pH 7.35). Hippocampal slices containing the anterior hippocampus (300 μm thick) were cut coronally in a Leica vibroslicer (VT1200 S; Leica Microsystems, Wetzlar, Germany) in the same cold solution. Slices were transferred to an incubation chamber with a nylon mesh containing aCSF2 (in mM, 119 NaCl, 1.25 NaH_2_PO_4_, 25 NaHCO_3_, 2.5 KCl, 2.5 CaCl_2_, 1.5 MgSO_4_, 11 glucose, purged with 95% O2/ 5% CO2, pH 7.35). Slices were kept at 37°C for 45-60 min for optimal recovery. After that, the incubation chamber was gently transferred out of the bath and held at room temperature (22–25 °C) for at least 1h before starting the recordings.

### Whole-cell recordings

The recording chamber consisted of a circular well of a 1–2 ml volume and was continuously perfused with aCSF2 at a flow rate of 4–5 ml/min. A horseshoe shape wire enchased with nylon wires was placed on top of the slice to allow for the most rapid flow while minimizing cell movement. The recording chamber was mounted on an upright fluorescence microscope (SliceScope Pro 1000, Scientifica). The microscope was used to identify individual cells from the CA1 or CA3 region of the hippocampus and, and after patching, spontaneous mini excitatory postsynaptic currents (mEPSCs) were recorded. Recordings were obtained in “gap free” model of 30 to 600 second recording intervals sampled at 10KHz and low-pass filtered at 1 KHz. Glass pipettes were pulled with a micropipette puller Model P-1000 (Sutter Instruments, USA) and had a resistance of 3–6 MΩ when filled with an internal solution (consisting of the following, in mM: 115 CsMeSO_3_, 20 CsCl, 10 HEPES, 2.5 MgCl_2_, 4 Na_2_-ATP, 0.4 Na-GTP, 10 Na-phosphocreatine, 0.6 EGTA, pH 7.2). Cells were voltage-clamped at −70 mV, and experiments were conducted only after the access resistance had stabilized. Membrane and access resistance were monitored before starting the recording and at the end of it. Recordings were included for analysis if there was less than a 20% variation in series resistance (15-35 MΩ) and the input resistance remained constant throughout the experiment (100–300 MΩ). 50 μM picrotoxin, 50 μM APV and 1 μM tetrodotoxin were added to the recording solution to avoid iPSCs contamination, NMDAR-mediated currents and EPSCs generated by synaptic transmission, respectively. All recordings were amplified and stored using amplifier Multiclamp 700B (Molecular Devices, San Jose, CA). Miniature events were detected and analysed with IGOR Pro 6.06 (Wavemetrics) using NeuroMatic 2.03 ((Rothman and Silver, 2018); http://www.neuromatic.thinkrandom.com). Statistical analysis was performed using GraphPad Prism version 8.0.1 for Mac OS X (GraphPad Software, San Diego California USA, www.graphpad.com).

The deactivation kinetics of AMPAR-mediated miniature responses were determined by fitting the average of the events for a given cell to a double-exponential function to calculate the weighted time constant (τ_w_):

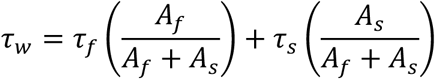

where A_f_ and τ_f_ are the amplitude and time constant of the fast component of recovery and A_s_ and τ_s_ are the amplitude and time constant of the slow component.

### Allen Brain Atlas RNA *in situ* hybridization (ISH) data analysis

Four different scientists manually inspected RNA *in situ* hybridization (ISH) data from adult mouse brain from the Allen Brain Atlas^94^. Each researcher reviewed the 283 proteins overexpressed in CA3-CA1, DG-CA3 and EC-DG synapses. RNA ISH data from the entorhinal cortex was also reviewed for proteins with highest expression in dDG. For a protein to be classified as with highest expression in one or more regions there had to be agreement on 3 out of the 4 researchers. Proteins were classified into those with concordant protein and ISH expression and non-concordant ones. Proteomic data was considered concordant with ISH data when the RNA expression level of a synaptic protein found with highest expression in one of the three hippocampal regions investigated had highest ISH levels in the somas of one or both brain regions contributing to that synapse. For instance, a protein found with highest expression in CA3-CA1 synapses had concordant ISH data if CA3 and/or CA1 somas presented highest expression level of that gene for 3 out of the 4 researchers.

### Pathway enrichment analysis

Pathway enrichment analysis was performed using the pathfindR R package^55^. pathfindR takes into consideration protein-protein interaction (PPI) data for pathway enrichment analysis, which is performed using one-sided hypergeometric tests. For our analysis PPI data was retrieved from BioGRID build 4.3.196 (https://thebiogrid.org/) and STRING version 11 (https://string-db.org/), both restricted to *Mus musculus* species. Only STRING interactions with a confidence score above 0.9 were taken into consideration. Redundant interactions between both databases were removed, resulting in a final interaction database with 339.776 interactions. Gene name conversions needed for merging data from different databases and converting them to updated gene symbols were done with biomaRt R package^95^. Pathways investigated with pathfindR were taken from MSigDB collections, (https://www.gsea-msigdb.org) and were restricted to *Mus musculus*. MSigDB contains several collections of gene sets, we used the C2 set: curated gene sets and the C5 set: ontology gene sets. On C2 collection, only REACTOME, WikiPathways and KEGG pathways were used for analysis, which resulted in 2405 gene sets. For the C5 collection all the GO gene sets were selected: Biological process (BP), Cellular Component (CC) and Molecular Function (MF), resulting in 10185 gene sets.

Briefly, pathfindR first builds a Protein Interacting Network (PIN) from all differentially expressed (DE) molecules (genes/proteins) investigated using the PPI data provided. Next, subnetworks are built from the PIN with a minimum length of 10 DE molecules using the Greedy algorithm with a maximum depth of 1, hence only considering the addition of direct neighbours from DE molecules. Subnetworks with 50% of gene overlap are discarded, maintaining those with a higher score, based on the adjusted p-value of DE molecules. Finally, pathway enrichment analyses is done for each subnetwork, using all the molecules of the PIN as the background set. Pathways that include less than 3 DE molecules are discarded. As the greedy algorithm is a stochastic method, the whole process is repeated 50 times, starting from the subnetwork construction. For a pathway to be considered it had to appear (occurrence) at least in 13 of the 50 (>25%) iterations. Finally, to reduce complexity, enriched pathways are grouped using hierarchical clustering, based on their similarity on the DE molecules they include. One ‘Representative’ term for each cluster was chosen based on the lowest p-value from the hypergeometric test. Heatmaps to represent gene/protein abundance data were generated with the scrattch.hicat R package from the Allen brain atlas (https://github.com/AllenInstitute/scrattch.hicat). Protein and RNA abundance data was normalized by a Log2(x+1) transformation and converted to z-scores.

Source data files relevant to these analyses: Source_Data_6, 7 and 8.

### Analysis of single cell RNA-sequencing data from the Allen Brain Cell Atlas

Single cell RNA-seq. data from mouse glutamatergic neurons of the hippocampal formation was retrieved from the Allen Brain Cell Atlas Database (Whole Cortex & Hippocampus - 10X Genomics (2020) with 10X-SMART-SEQ taxonomy^49^). More precisely, we collected RNA-seq. data from the following sub-classes of glutamatergic neurons: DG, CA2-IG-FC, CA3, CA1-ProS, SUB-ProS, CT SUB and NP SUB, all belonging to the hippocampal formation which also includes subiculum neurons^49^. Of note, in this manuscript we refer to ABA Sub-classes as Classes, for simplicity.

Statistical analysis of RNA abundance data was performed using the Seurat R package^96^, which is designed to work with single cell gene expression data. To identify DE genes, we performed the Wilcoxon Rank Sum test, which is the default test in the Seurat package. p-values were corrected for multiple testing using the Benjamini-Hochberg procedure. As we are interested in identifying abundance differences among genes expressed at synapses, we only worked with RNA abundance data from the genes corresponding to our reference list of synaptic proteins (Suppl. Table 1).

To identify DE genes in each group (i.e. class or type) we compared gene expression in that group against that of all other groups together. The identification of DE among neuronal types was done within classes. Statistics were done with an equal number of neurons for each group. To identify DE genes between classes we used 100 neurons per class, and to identify DE genes between neuronal types we used 25 neurons per type. To sample a representative number of neurons per group so that all DE genes per group would be identified we had to iterate this process. We empirically found that 150 iterations were enough to saturate the number of DE genes in each group. Importantly, for a gene be considered as DE in each group it had to be identified as significantly DE in at least 90% of these 150 iterations. Furthermore, DE genes not only had to present and adjusted p-value below 0.05, but their expression fold change value (in log2 scale) had to be above 0.6 for overexpressed genes or below −0.6 for downregulated genes.

Gene expression dendograms were generated with the median value of log2(x+1) transformed gene expression abundance data and using the scrattch.hicat R package from the Allen brain atlas (https://github.com/AllenInstitute/scrattch.hicat).

Source data files relevant to these analyses: Source_Data_1 to 5.

### General Statistics

Specific statistical tests are mentioned in the figure legends. Data was tested for normality using the Shapiro-Wilk test and the Kolmogorov-Smirnov test. When possible, statistical test used were two-sided. Statistical tests on omics data were corrected for multiple testing.

### Permutation test

A permutation test was performed to assess whether the observed concordance between protein and RNA abundance was higher than expected by chance. To calculate the expected concordance, differentially expressed (DE) proteins were randomly assigned to the three synaptic types, and the concordance with the ABCA RNA abundance data was computed. This process was repeated 1,000 times, yielding an average concordance which would correspond to random or chance concordance. Finally, the permutation test, based on these 1,000 random sets, computes if the observed concordance iss significantly different from the one expected by chance. Permutation test was performed with the function permutation_test included in the Python package scipy.stats.

### Uniform Manifold Approximation and Projection (U-MAPS)

To generate neuronal classes and types, gene expression U-MAPS we used the umap-learn package (https://pypi.org)^97^. The hyperparameters used to generate the maps were: Random state: 24, Number of neighbours: 15 and Minimum Distance 0.1. All other parameters were left as by default. Only the first two dimensions were used to generate the u-maps.

### Gene classification using machine learning

We used the random forest classification method to identify genes with the highest weight in the organization of neurons in classes and types. Gene expression data from the Allen Brain atlas was analysed with the ‘Random Forest Classifier’ function within the scikit-learn (https://scikit-learn.org/0.16/about.html) Python package^78^. The hyperparameters used for the Random Forest Classifier were: Random state: 24, Max. Depth: 12 and Number of estimators: 200. Values for all other parameters were kept as by default. The test set used included 20% of neurons in each group and the train set the remaining 80%. The ‘confusion matrix’ function from scikit-learn was used to generate confusion matrices.

Source data file relevant to these analyses: Source_Data_9.

## Source Data

Source_Data_1_Iteration_Classes.R: R script to iterate the statistical analysis performed with Seurat to identify genes differentially expressed between neuronal classes.

Source_Data_2_Iteration_Types.R: R script to iterate the statistical analysis performed with Seurat to identify genes differentially expressed between neuronal types.

Source_Data_3_Analysis_Classes.R: R script to generate data tables and graphs for genes differentially expressed between neuronal classes. This script also includes a quality control test to validate differentially expressed genes.

Source_Data_4_Analysis_Types.R: R script to generate data tables and graphs for genes differentially expressed between neuronal Types. This script also includes a quality control test to validate differentially expressed genes.

Source_Data_5_Split_Types.R: R script to obtained data from a subset of neuronal types from the entire transcriptomic database provided by the ABCA.

Source_Data_6_pathfindR_Proteomics.Rmd: R script to perform the pathfinder analysis and to generate the heatmaps from the proteomics data.

Source_Data_7_pathfindR_Classes.R: R script to perform the pathfinder analysis and to generate the heatmaps from transcriptomics data of neuronal classes (ABCA).

Source_Data_8_pathfindR_Types.R: R script to perform the pathfinder analysis and to generate the heatmaps from transcriptomics data of neuronal types (ABCA).

Source_Data_9_ Random_Forest.ipynb: Python code to perform the Random Forest analysis on transcriptomic data from the ABCA.

## Data and code Availability

All the data generated by the bioinformatics analysis performed in this manuscript can be found in the supplementary tables.

Mass spectrometry proteomics data has been deposited to the ProteomeXchange Consortium via the PRIDE partner repository^98^ with the dataset identifiers PXD052901 and PXD052913.

All custom-made code is available from GitHub: https://github.com/Alex-Bayes/Synaptic-Proteome-Diversity

## Acknowledgments

RRV, DdCB, OZR and AB financial support was provided by: PID2021-124411OB-I00 and RTI2018-097037-B-I00 (MINECO/MCI/AEI/FEDER, EU), Award AC17/00005 by ISCIII through AES2017 and within the NEURON framework, Ramón y Cajal Fellowship (RYC-2011-08391p), IEDI-2017-00822, Universitat Autònoma de Barcelona and AGAUR (2017 SGR 1776 and 2021 SGR 01005). DdCB thanks AGAUR/Generalitat de Catalunya/FEDER, EU for ‘Ajuts per a la contractació de personal investigador novel (FI)’ Ref.2020FI_B00130. All authors thank the CERCA Programme/Generalitat de Catalunya for institutional support. JP, AC and DS were supported by grant from Ministerio de Universidades, Ciencia y Innovación Innovación (PID2020-119932GB-I00; MCIN/ AEI /10.13039/5011000011033) and the María de Maeztu (MDM-2017-0729 to Institut de Neurociencies, Universitat de Barcelona).

## Author Contributions

RRV, DdCB, ABP, DAA, OZR, JP, AC and DRV performed experiments. NR, DS, CS and AB designed and supervised all experiments and secured funding. AB and CS wrote the manuscript. All authors reviewed and approved the manuscript.

## Competing interests

Authors declare no competing interests.

## Notes

### Competing Interest Statement

The authors have declared no competing interest.

### Summary of Updates

We have updated the following sections: Title, Abstract, Results, Discussion and Methods. We include new experiments: Double Immunofluorescence (Fig.2 and Suppl. Figs. 4 to 6) Whole cell recording of mEPSCs (Suppl Fig. 7) We have updated the authors list.

