## Supplementary Information for "Synaptic proteome diversity is shaped by the levels of glutamate receptors and their regulatory proteins"

### Supplementary Figures

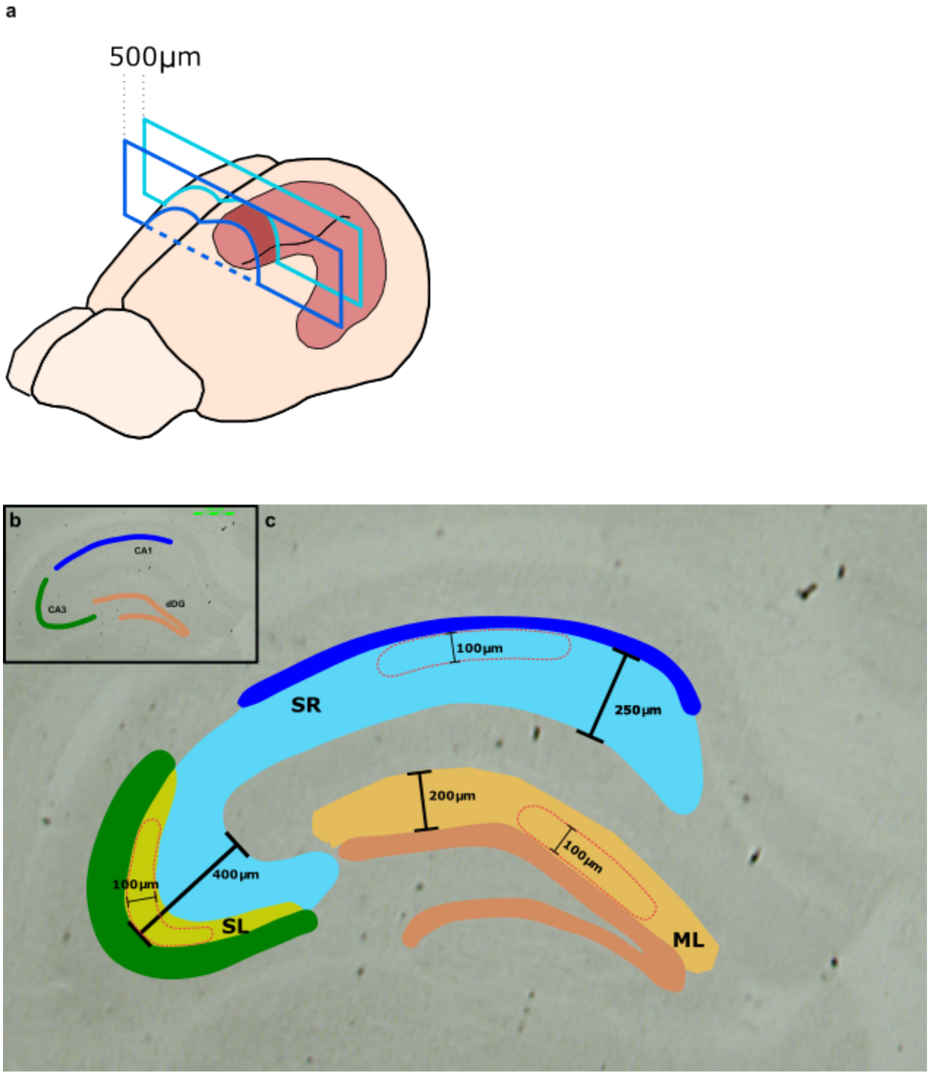


**Supplementary Figure 1. Strategy used to microdissect the hippocampal layers containing the synapses of the trisynaptic circuit.**

**a.** Drawing of a mouse brain showing the localization of the hippocampus, in red. The portion of the dorsal hippocampus that was analysed in this work is shown in dark red. This covered approximately 500 µm in the longitudinal axis of the brain.

**b.** Brightfield image showing the hippocampus in a coronal section of the dorsal mouse brain. The 3 subfields, CA1, CA3 and dDG, investigated in this study are indicated. CA1 and CA3 pyramidal layers and DG granular layer are shown differently coloured. Scale bar 1000µm.

**c.** Anatomical localization and dimensions, particularly width, of the different hippocampal layers from which we collected microdissected neuropile. Pyramidal and granular layers coloured as in (a). Shapes delimited by red dashed lines represent examples of microdissected neuropile fragments, where their width is also shown. Collected fragments in all subfields had 100µm in width, approximately. Neuropile fragments were collected from the following layers: i) Stratum Radiatum (SR, in blue) in the CA1 subfield, ii) Stratum Lucidum (SL, in pale green) in the CA3 subfield and iii) Molecular Layer (ML) at the dorsal Dentate Gyrus.

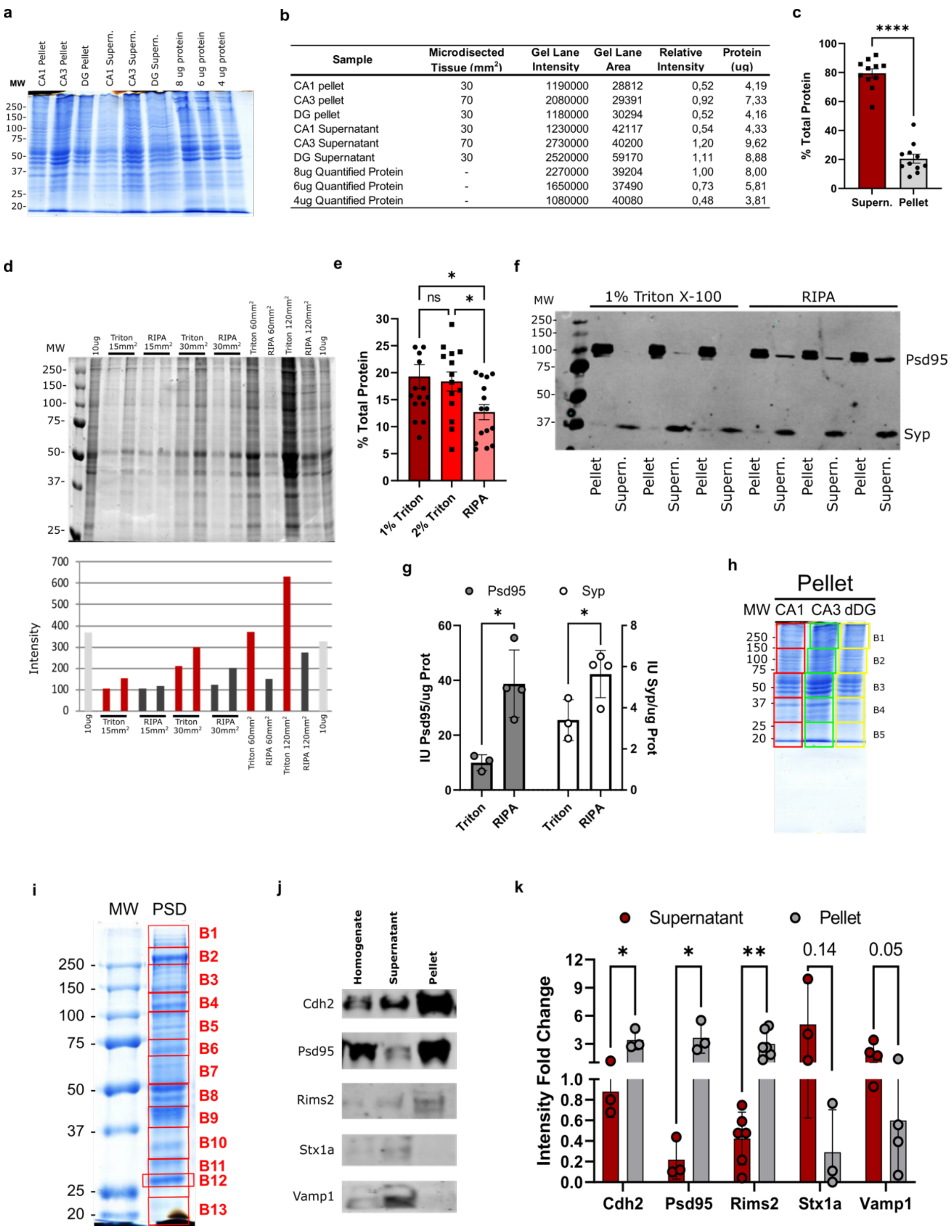


| Sample | Microdisected Tissue (mm²) | Gel Lane Intensity | Gel Lane Area | Relative Intensity | Protein (ug) |
|---|---|---|---|---|---|
| CA1 pellet | 30 | 1190000 | 28812 | 0,52 | 4,19 |
| CA3 pellet | 70 | 2080000 | 29391 | 0,92 | 7,33 |
| DG pellet | 30 | 1180000 | 30294 | 0,52 | 4,16 |
| CA1 Supernatant | 30 | 1230000 | 42117 | 0,54 | 4,33 |
| CA3 Supernatant | 70 | 2730000 | 40200 | 1,20 | 9,62 |
| DG Supernatant | 30 | 2520000 | 59170 | 1,11 | 8,88 |
| 8ug Quantified Protein | - | 2270000 | 39204 | 1,00 | 8,00 |
| 6ug Quantified Protein | - | 1650000 | 37490 | 0,73 | 5,81 |
| 4ug Quantified Protein | - | 1080000 | 40080 | 0,48 | 3,81 |

**Supplementary Figure 2. Optimization and validation of the biochemical procedure used to obtain preparations enriched in synaptic structures and proteins.**

**a.** Protein gel electrophoresis stained with Coomassie Brilliant Blue with 1% Triton X-100 insoluble (Pellet) and soluble (Supern.) fractions obtained from microdissected neuropile of the three hippocampal subfields. 8, 6 and 4 μg of precisely quantified protein from hippocampal synaptic fractions isolated by density gradient ultracentrifugation were used to estimate protein abundance of fractions derived from microdissected neuropile.

**b.** Quantification of signal intensity of gel lanes in (a) and estimation of protein abundance in biochemical fractions from microdissected neuropile.

**c.** Percentage of total protein from microdissected neuropile recovered in 1% Triton X-100 soluble (Supern.) and insoluble (Pellet) fractions. Error bars: SE. Sample size (n) = 3, 3 replicates per sample. Statistics, unpaired two-sided Student's T-Test, **** < 0.0001.

**d.** **Top:** Silver-stained protein gel with biochemical fractions insoluble to 1% Triton X-100 or RIPA buffer obtained from increasing areas ($mm^2$) of microdissected neuropile. 10μg of quantified protein from hippocampal synaptic fractions isolated by density gradient ultracentrifugation were added to the gel for reference. **Bottom:** Bar chart with signal intensity from gel lanes in above image.

**e.** Percentage of protein recovered in the pellet fractions of microdissected tissue treated with a buffer containing 1% Triton X-100 (dark red column), 2% Triton X-100 (red column) or RIPA buffer (light red column). Error bars: SE. Sample size (n) = 4, 3-4 replicates per sample. Statistics, One-way ANOVA and Fisher's LSD post-hoc test, * $p < 0.05$.

**f.** Immunoblot of Triton and RIPA insoluble (Pellet) and soluble (Supern.) fractions obtained from hippocampal microdissected neuropile. Proteins investigated are Psd95, mostly insoluble to triton and Synaptophysin (Syp) a synaptic vesicle protein, mostly soluble to triton.

**g.** Bar plot of Psd95 (grey bars) and Synaptophysin (Syp, white bars) abundance as determined by immunoblot of 1% Triton X-100 soluble fractions from microdissected hippocampal neuropile. Error bars: SE. Sample size (n) = 4. Statistics, unpaired two-sided Student's T-test, * $p < 0.05$.

**h.** Protein gel electrophoresis of 1% Triton X-100 insoluble pellets stained with Coomassie Brilliant Blue. 5 gel bands (B1-B5) were collected and processed independently in the proteomics workflow.

**i.** Protein gel electrophoresis of synaptic preparations isolated by density gradient ultracentrifugation from mouse hippocampi stained with Coomassie Brilliant Blue. 13 bands (B1-B13) were collected and processed independently in the proteomics workflow to generate a synaptic reference proteome.

**j.** Immunoblots of a transsynaptic protein involved in cell adhesion (Cdh2, cadherin 2), a postsynaptic scaffolding molecule (Psd95), and three presynaptic proteins located at: i) active zone (Rims2), SNARE complex (Stx1a) and synaptic vesicles (Vamp1). Samples analysed are whole extract (Homogenate) and triton soluble (Supernatant) and insoluble (Pellet) fractions from microdissected hippocampal neuropile.

**k.** Abundance of proteins in (j) relative to their abundance in the homogenate fraction. Statistically significant difference in protein abundance between the supernatant and pellet fractions is indicated. Error bars: SE. Sample size (n) = 6. Statistics, unpaired two-sided Student's T-test, * $p < 0.05$, ** $p < 0.01$.

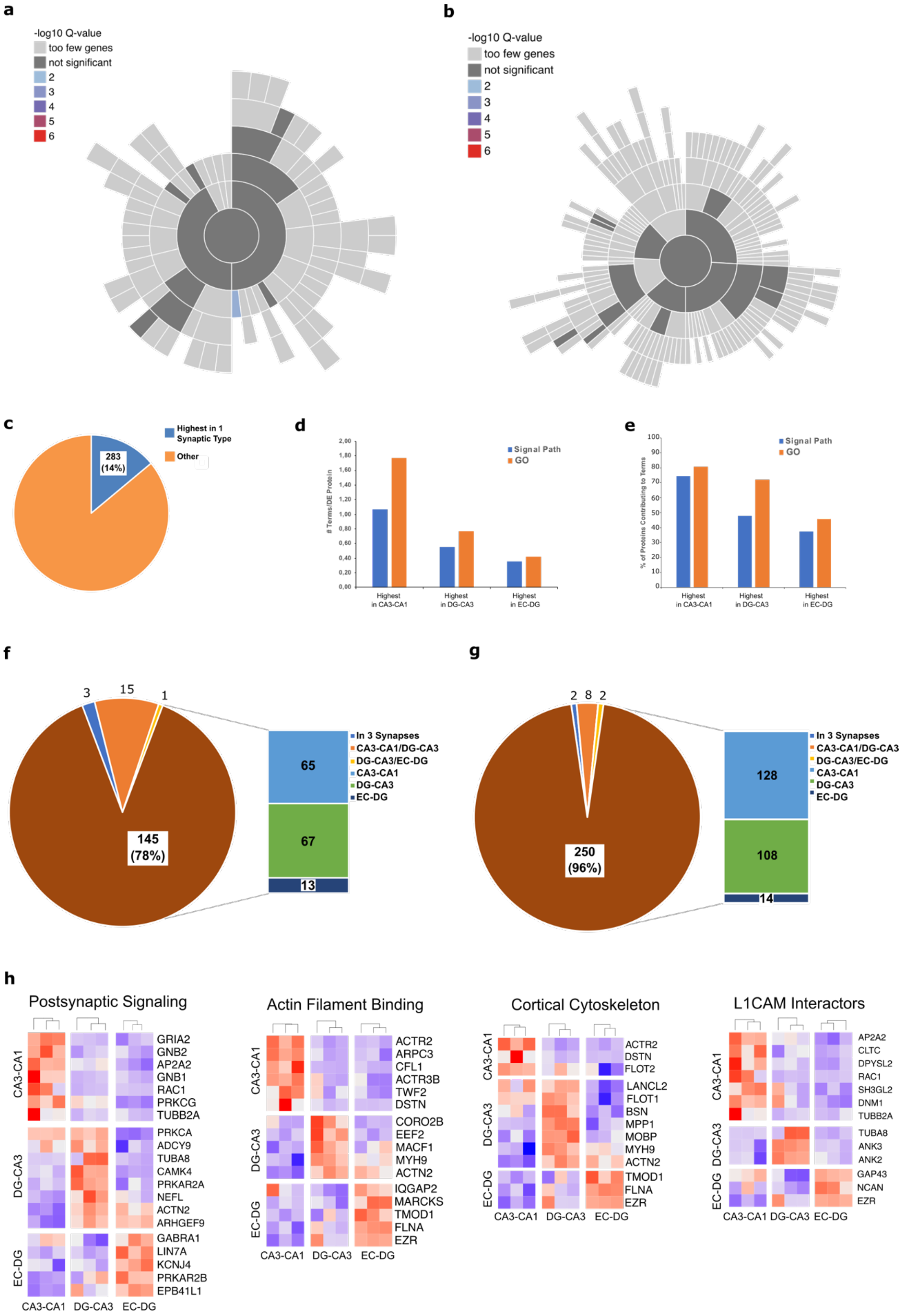


**Supplementary Figure 3. Most Signalling Pathways and Gene Ontology (GO) terms found in proteins differentially expressed between synaptic types are synapse-specific**

**a.** Sunburst plot showing SynGO Cellular Component terms enriched among proteins removed after benchmarking with the reference synaptic proteome. Note that not only one term (postsynaptic ribosome, adj. p-value 4.4e-5) is significantly overrepresented among this protein set.

**b.** Sunburst plot showing SynGO Biological Process terms enriched among proteins removed after benchmarking with the reference synaptic proteome. No terms appear significantly overrepresented.

**c.** Proportion of proteins identified by proteomics with a statistically significant highest expression in one of the three synaptic types investigated.

**d.** Ratio of significantly overrepresented terms per differentially expressed protein in each sample investigated. Signalling pathways (blue bars) investigated are from the databases Reactome, KEGG and Wikipathways. Gene ontology (GO, orange bars) terms investigated belong to the domains Cellular Component, Biological Process and Molecular Function.

**e.** Percentage of differentially expressed proteins contributing to signalling pathways (blue bars) and GO terms (orange bars).

**f.** Number of signalling pathways significantly overrepresented in 3, 2 or 1 hippocampal layers.

**g.** Number of significantly overrepresented GO terms among proteins with highest expression in 3, 2 or 1 synapses.

**h.** Heatmaps showing relative protein abundance in the 9 samples analysed by proteomics, three biological replicates per hippocampal layer. High abundance shown in red and low in blue. Abundance of proteins in four pathways/terms found significantly enriched in all three synapses is presented.

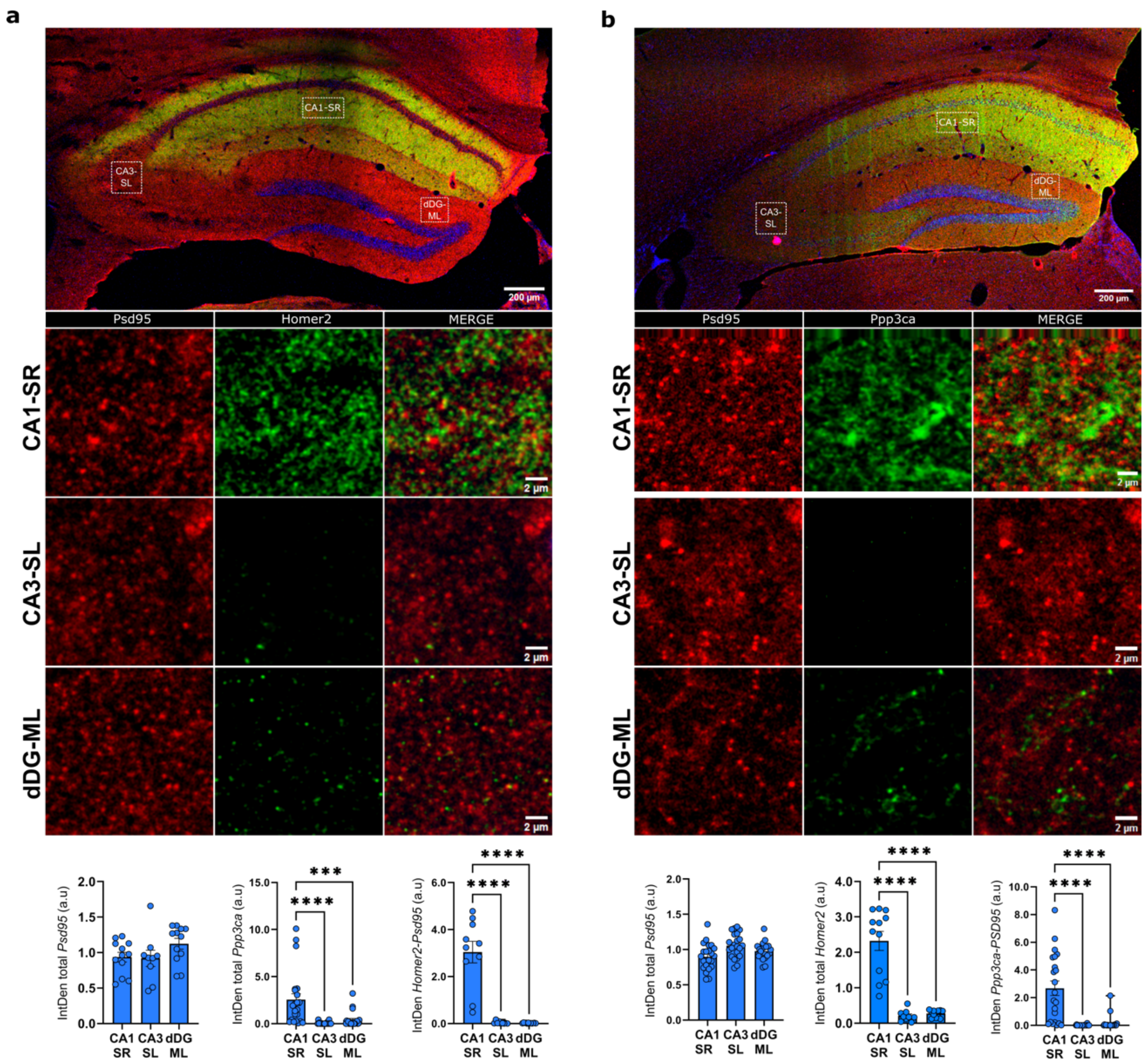


**Supplementary Figure 4. Validation by double immunofluorescence of proteins found differentially expressed at CA3-CA1 synapses.**

**a.** Increased expression of Homer2 in CA3-CA1 synapses at the stratum radiatum.

**Top image**: Complete hippocampus. Red signal corresponds to Psd95 and green signal to Homer2. White doted shapes denote locations where high-magnification images were taken in CA1 - stratum radiatum (CA1-SR), in CA3 - stratum lucidum (CA3-SL) and in Dentate Gyrus - dorsal molecular layer (dDG-ML). Scale bar 200 µm.

**High magnification images**: columns correspond to Psd95, Homer2 and the merged signal. Rows correspond to images from CA1-SR, CA3-SL and dDG-ML. Scale bars 2 µm.

**Bottom:** Bar and dots plots show the quantification of total mean fluorescence intensity of Psd95 (left), total mean fluorescence intensity of Homer2 (middle) and mean fluorescence intensity of the overlapping signal (right). Error bars: SE. Sample size (n) = 4, 3-6 images taken from each subfield and animal. Statistics, one-way ANOVA and Fisher's LSD post-hoc test, (*** $p < 0.001$, **** $p < 0.0001$).

**b.** Increased expression of Calcineurin (Ppp3ca) in CA3-CA1 synapses at the stratum radiatum.

**Top image**: Complete hippocampus. Red signal corresponds to Psd95 and green signal to Ppp3ca. White doted shapes denote locations where high-magnification images were taken in CA1 - stratum radiatum (CA1-SR), in CA3 - stratum lucidum (CA3-SL) and in Dentate Gyrus - dorsal molecular layer (dDG-ML). Scale bar 200 µm.

**High magnification images**: columns correspond to Psd95, Ppp3ca and the merged signal. Rows correspond to images from CA1-SR, CA3-SL and dDG-ML. Scale bars 2 µm.

**Bottom:** Bar and dots plots show the quantification of total mean fluorescence intensity of Psd95 (left), total mean fluorescence intensity of Ppp3ca (middle) and mean fluorescence intensity of the overlapping signal (right). Error bars: SE. Sample size (n) = 4, 3-6 images taken from each subfield and animal. Statistics, one-way ANOVA and Fisher's LSD post-hoc test, (**** $p < 0.0001$).

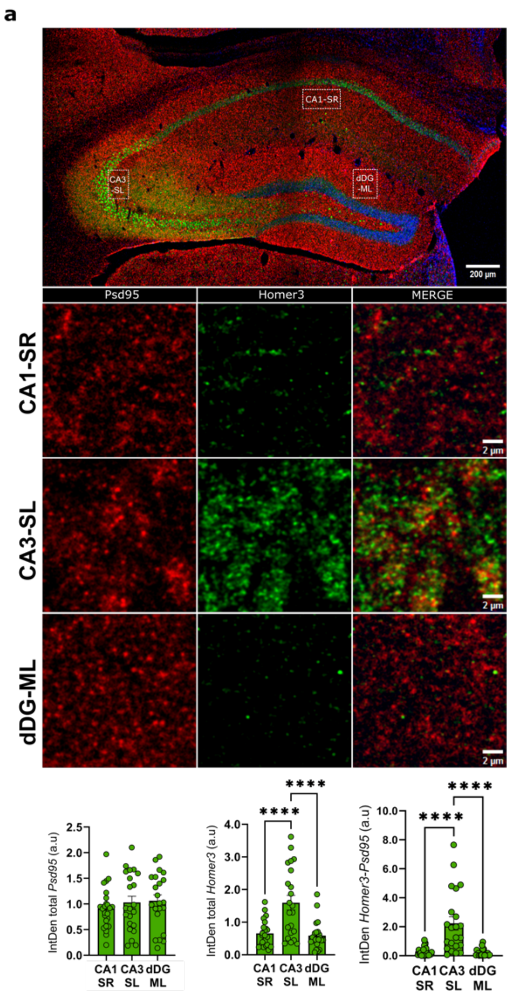


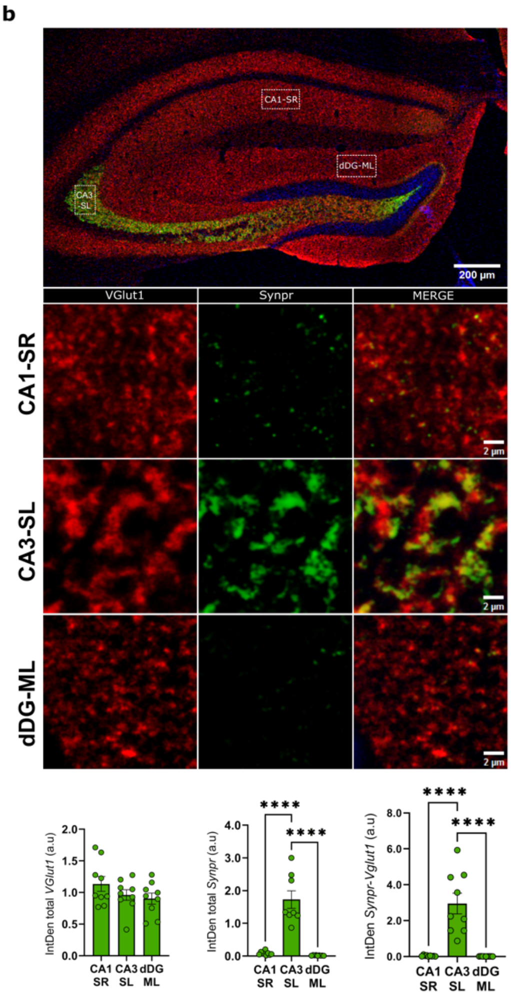


**Supplementary Figure 5. Validation by double immunofluorescence of proteins found differentially expressed at DG-CA3 synapses.**

**a.** Increased expression of Homer3 in DG-CA3 synapses at the stratum lucidum.

**Top image**: Complete hippocampus. Red signal corresponds to Psd95 and green signal to Homer3. White doted shapes denote locations where high-magnification images were taken in CA1 - stratum radiatum (CA1-SR), in CA3 - stratum lucidum (CA3-SL) and in Dentate Gyrus - dorsal molecular layer (dDG-ML). Scale bar 200 µm.

**High magnification images**: columns correspond to Psd95, Homer3 and the merged signal. Rows correspond to images from CA1-SR, CA3-SL and dDG-ML. Scale bars 2 µm.

**Bottom:** Bar and dots plots show the quantification of total mean fluorescence intensity of Psd95 (left), total mean fluorescence intensity of Homer3 (middle) and mean fluorescence intensity of the overlapping signal (right). Error bars: SE. Sample size (n) = 4, 3-6 images taken from each subfield and animal. Statistics, one-way ANOVA and Fisher's LSD post-hoc test, (**** $p < 0.0001$).

**b.** Increased expression of Synaptoporin (Synpr) in CA3-CA1 synapses at the stratum lucidum.

**Top image**: Complete hippocampus. Red signal corresponds to vGlut1 and green signal to Ppp3ca. White doted shapes denote locations where high-magnification images were taken in CA1 - stratum radiatum (CA1-SR), in CA3 - stratum lucidum (CA3-SL) and in Dentate Gyrus - dorsal molecular layer (dDG-ML). Scale bar 200 µm.

**High magnification images**: columns correspond to vGlut1, Ppp3ca and the merged signal. Rows correspond to signal from CA1-SR, CA3-SL and dDG-ML. Scale bars 2 µm.

**Bottom:** Bar and dots plots show the quantification of total mean fluorescence intensity of vGlut1 (left), total mean fluorescence intensity of Synpr (middle) and mean fluorescence intensity of the overlapping signal (right). Error bars: SE. Sample size (n) = 4, 3-6 images taken from each subfield and animal. Statistics, one-way ANOVA and Fisher's LSD post-hoc test, (**** $p < 0.0001$).

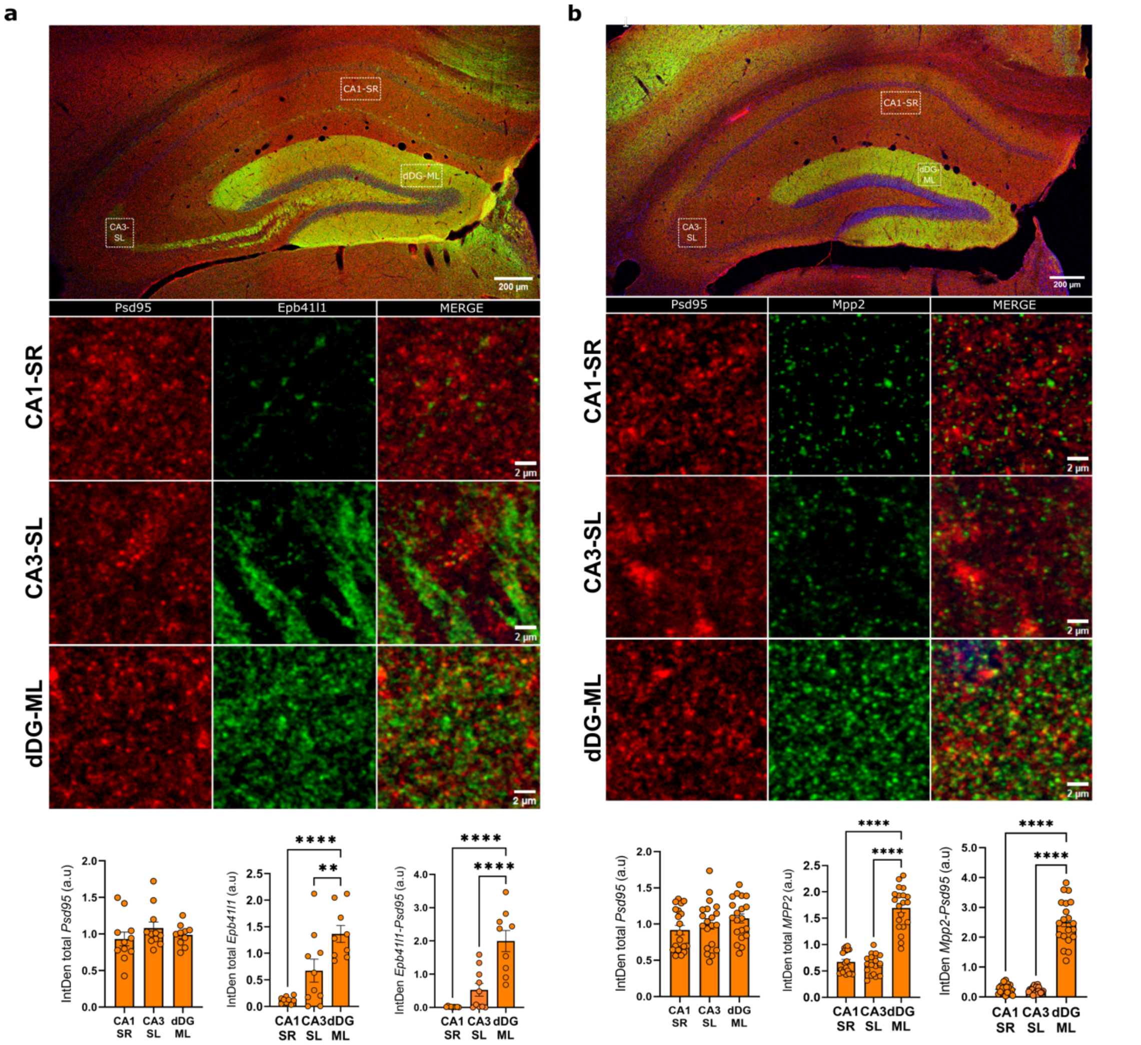


**Supplementary Figure 6. Validation by double immunofluorescence of proteins found differentially expressed at EC-DG synapses.**

**a.** Increased expression of Band 4.1-like protein 1 (Epb41l1) in EC-DG synapses at the dorsal molecular layer.

**Top image**: Complete hippocampus. Red signal corresponds to Psd95 and green signal to Epb41l1. White doted shapes denote locations where high-magnification images were taken in CA1 - stratum radiatum (CA1-SR), in CA3 - stratum lucidum (CA3-SL) and in Dentate Gyrus - dorsal molecular layer (dDG-ML). Scale bar 200 µm.

**High magnification images**: columns correspond to Psd95, Epb41l1 and the merged signal. Rows correspond to images from CA1-SR, CA3-SL and dDG-ML. Scale bars 2 µm.

**Bottom:** Bar and dots plots show the quantification of total mean fluorescence intensity of Psd95 (left), total mean fluorescence intensity of Epb41l1 (middle) and mean fluorescence intensity of the overlapping signal (right). Error bars: SE. Sample size (n) = 4, 3-6 images taken from each subfield and animal. Statistics, one-way ANOVA and Fisher's LSD post-hoc test, (** $p < 0.01$, **** $p < 0.0001$).

**b.** Increased expression of Mpp2 in EC-DG synapses at the dorsal molecular layer.

**Top image**: Complete hippocampus. Red signal corresponds to Psd95 and green signal to Mpp2. White doted shapes denote locations where high-magnification images were taken in CA1 - stratum radiatum (CA1-SR), in CA3 - stratum lucidum (CA3-SL) and in Dentate Gyrus - dorsal molecular layer (dDG-ML). Scale bar 200 µm.

**High magnification images**: columns correspond to Psd95, Mpp2 and the merged signal. Rows correspond to images from CA1-SR, CA3-SL and dDG-ML. Scale bars 2 µm.

**Bottom:** Bar and dots plots show the quantification of total mean fluorescence intensity of Psd95 (left), total mean fluorescence intensity of Mpp2 (middle) and mean fluorescence intensity of the overlapping signal (right). Error bars: SE. Sample size (n) = 4, 3-6 images taken from each subfield and animal. Statistics, one-way ANOVA and Fisher's LSD post-hoc test, (**** $p < 0.0001$).

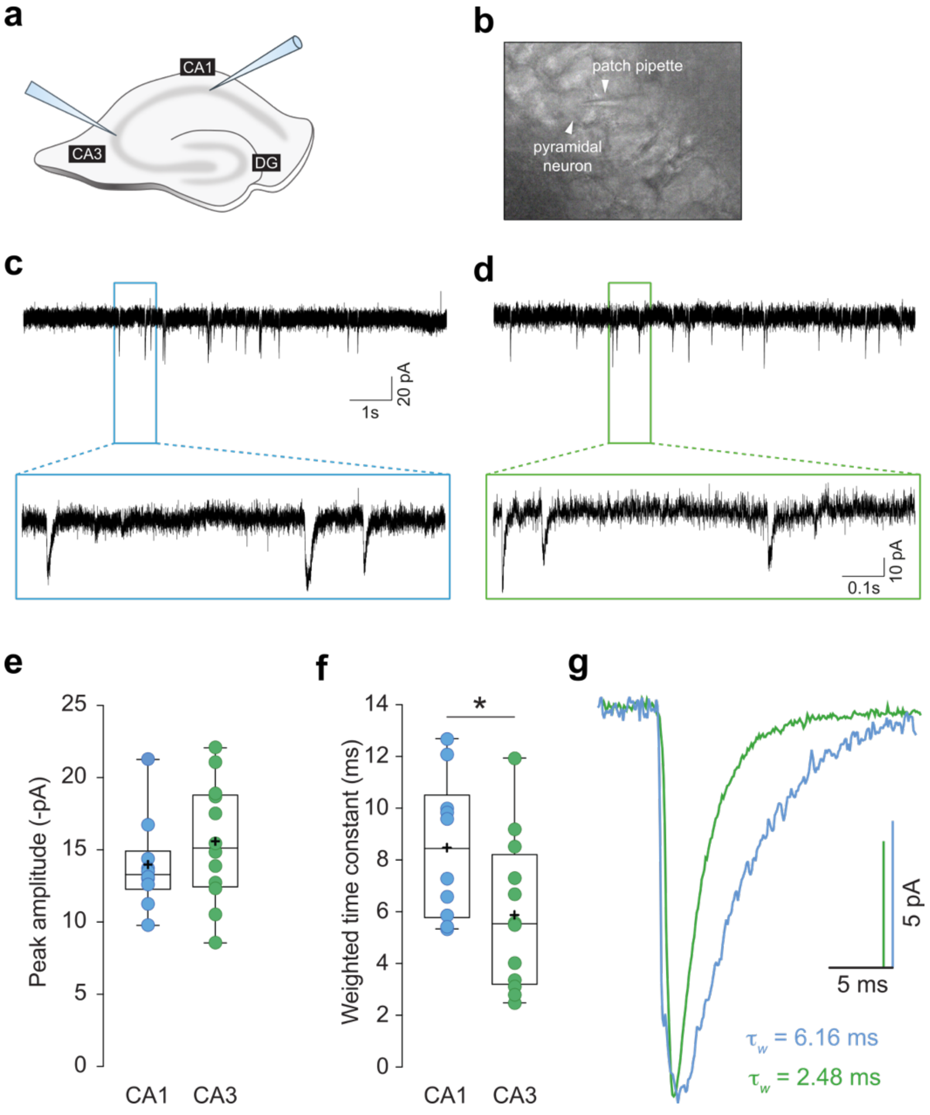


**Supplementary Figure 7. Different kinetics of AMPAR-mediated mEPSCs in CA1 and CA3 subfields indicate higher proportion of GluA2-containing AMPARs in the CA1 subfield.**

Miniature events driven by AMPAR activation were recorded from anterior coronal hippocampal slices.

**a.** Schematic representation of the preparation of anterior coronal hippocampal slices used for the recording of CA1 and CA3 individual pyramidal neurons. Patch pipettes indicate the regions where cells were recorded.

**b.** Example image of a pipette on a hippocampal pyramidal neuron from CA3 during a whole cell recording.

**c-d.** Representative traces recorded in CA1 (c) or CA3 (d) pyramidal neurons. Bottom panels show the recording with magnification for easier comparison of individual mEPSCs.

**e.** mEPSCs peak amplitude of CA1 (blue) and CA3 (green) pyramidal neurons. No statistical difference observed (Student's T-test). Error bars: SE. Sample size (n) = 12.

**f.** Weighted time constant of CA1 (blue) and CA3 (green) pyramidal neurons. Error bars: SE. Sample size (n) = 12. Unpaired two-sided Student's T-test; *$p<0.05$.

**g.** Peak-scaled average mEPSCs from a CA1(blue) and CA3 (green) hippocampal pyramidal neurons (average of 338 and 2735 traces for CA1 and CA3 recordings, respectively).

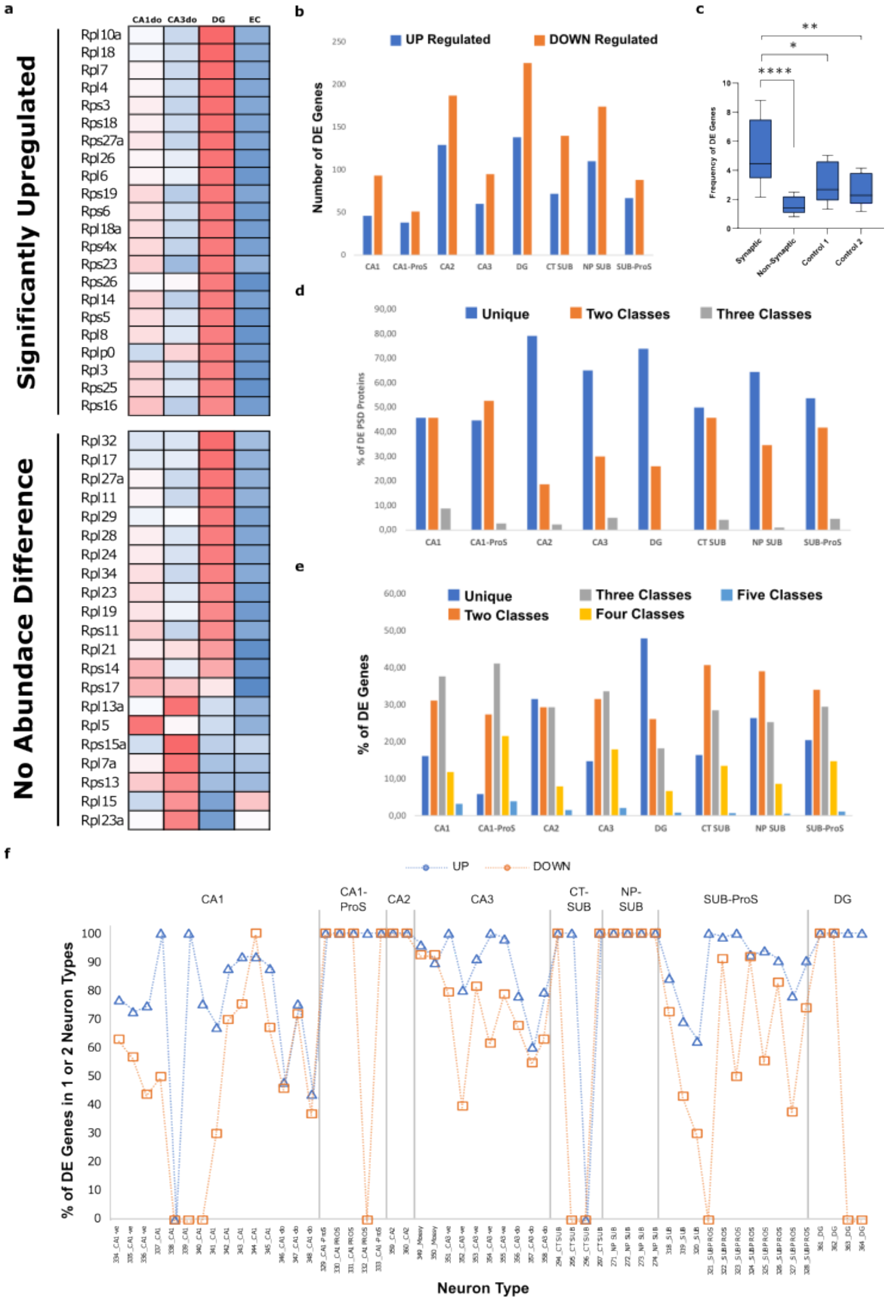


**Supplementary Figure 8. Significantly up-regulated genes are more specific to neuronal classes and types than down-regulated ones.**

**a.** Heatmap presenting normalized (z-score) mean RNA abundance of genes coding for ribosomal proteins from the 4 classes of excitatory neurons that constitute the trisynaptic loop in the dorsal (do) hippocampus (CA1do, CA3do, dentate gyrus -DG- and entorhinal cortex -EC-). All ribosomal gens showing statistically significant RNA expression differences were upregulated in the DG (top panel). Many genes for which expression differences did not reach statistical significance also display a tendency for increased expression in the DG (bottom panel). RNA expression data obtained from the Allen Brain Cell Atlas. Statistical analysis of RNA expression differences between neuronal classes was performed with the Seurat R package and the Wilcoxon Rank Sum test. Abundance scale, 2 (dark red) to -2 (dark blue).

**b.** Number of genes expressed at synapses found significantly up- (blue bars) or down- regulated (orange bars) in classes of excitatory neurons from the hippocampal formation.

**c.** Frequency of differentially expressed genes among different gene sets. Including genes expressed at synapses (synaptic), genes not expressed at synapses (Non-Synaptic), a random set of all genes of the same size of the synaptic set (Control 1) and a random set of non-synaptic genes of the same size of the synaptic set (Control 2). Statistics, Chi square Test, **** $p < 0.0001$, ** $p < 0.01$ and * $p < 0.05$.

**d.** Percentage of gens localized to synapses that are upregulated in one (blue), two (orange), or three (grey) classes of excitatory neurons.

**e.** Percentage of gens localized to synapses that are downregulated in one (blue), two (orange), three (grey), four (yellow) or five (light blue) classes of excitatory neurons.

**f.** Percentage of genes expressed at synapses being up-regulated (blue line) or down-regulated (orange line) in 1 or 2 excitatory neuron types from the hippocampal formation.

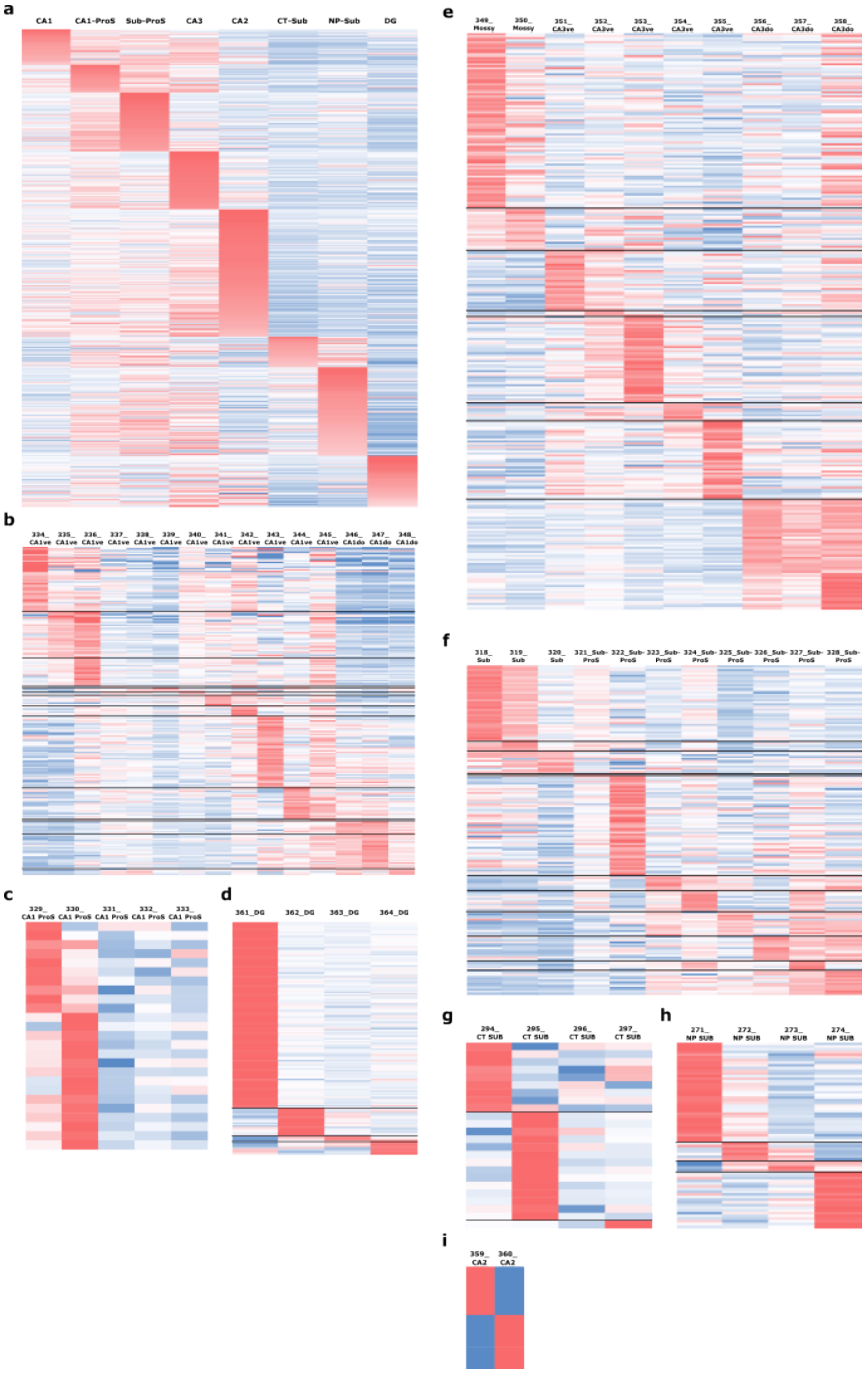


**Supplementary Figure 9. Gene expression heatmaps for synaptic genes expressed in neuronal classes and types.**

**a.** Heatmap showing relative RNA abundance data across all excitatory neuronal classes for genes found upregulated in each class.

**b.** Heatmap showing relative RNA abundance data across types of CA1 excitatory neurons for genes found upregulated in each type.

**c.** Heatmap showing relative RNA abundance data across types of CA1-ProS excitatory neurons for genes found upregulated in each type.

**d.** Heatmap showing relative RNA abundance data across types of dentate gyrus (DG) excitatory neurons for genes found upregulated in each type.

**e.** Heatmap showing relative RNA abundance data across types of CA3 excitatory neurons for genes found upregulated in each type.

**f.** Heatmap showing relative RNA abundance data across types of SUB-ProS excitatory neurons for genes found upregulated in each type.

**g.** Heatmap showing relative RNA abundance data across types of CT-SUB excitatory neurons for genes found upregulated in each type.

**h.** Heatmap showing relative RNA abundance data across types of NP-SUB excitatory neurons for genes found upregulated in each type.

**i.** Heatmap showing relative RNA abundance data across the two types of CA2 excitatory neurons for genes found upregulated in each type.

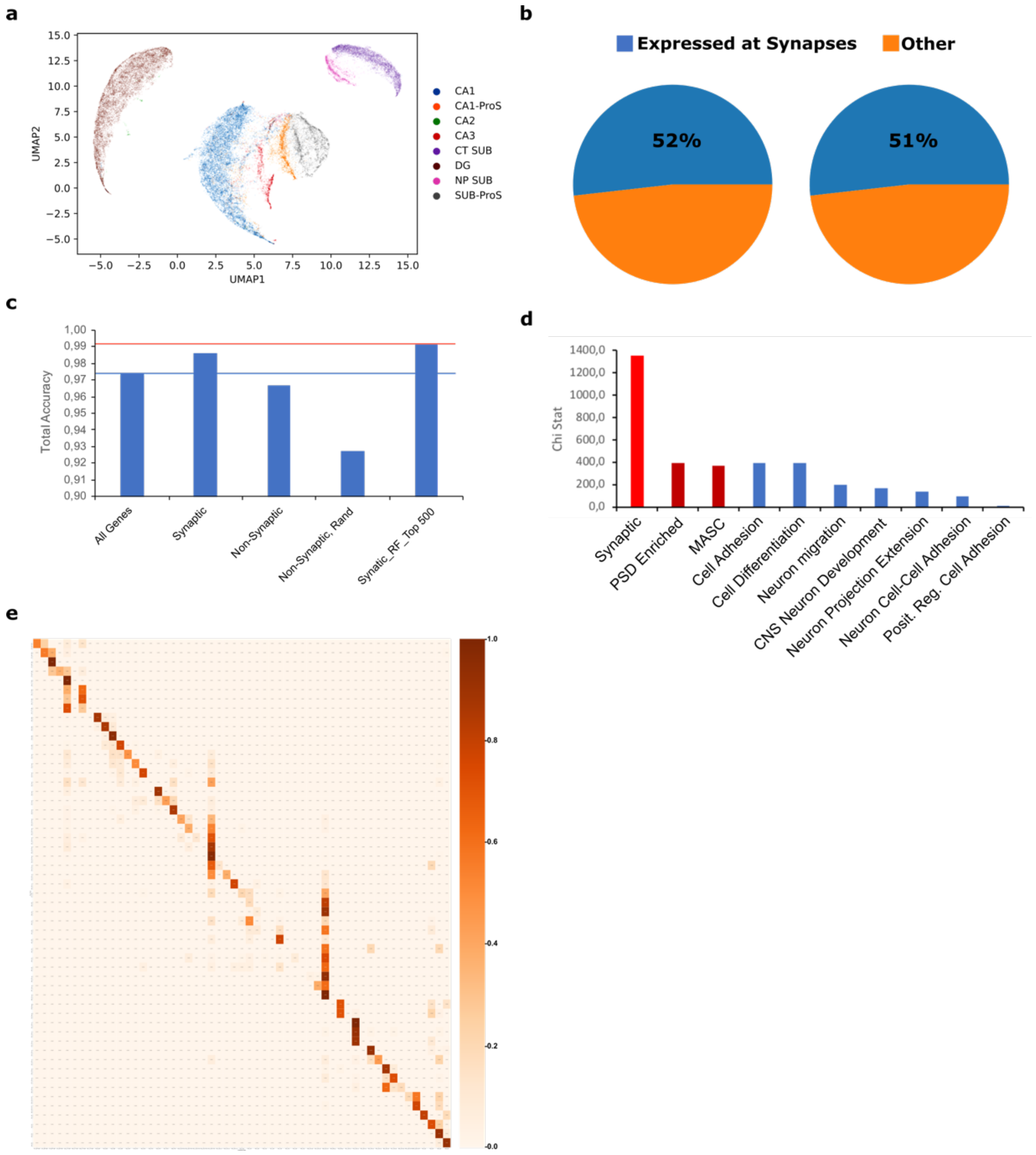


**Supplementary Figure 10. Synaptic proteins are highly relevant for the classification of excitatory neurons into classes and types.**

**a.** UMAP graph generated with scRNAseq data from genes not expressed at synapses from excitatory neurons in the 8 classes from the hippocampal formation.

**b.** Proportion of genes with synaptic (blue) or non-synaptic (orange) localization among the top 1000 genes contributing to the classification of neuronal classes (left chart) or neuronal types (right chart).

**c.** Bar plot showing total accuracy in neuronal class prediction determined by the Random Forest machine learning method using different gene sets to train the algorithm: i) all gens in the dataset, ii) genes with a synaptic localization, iii) genes with a non-synaptic localization, iv) a set of randomly selected genes not found at the synapse and of the same size of the set of genes localized at synapses and v) the set of 520 genes expressed at synapses among the 1000 genes mostly contributing to neuronal classification.

**d.** Bar plat with the Chi square statistic obtained from Chi square tests of overrepresentation of different gene sets among the 1000 genes mostly contributing to neuronal classification.

**e.** Confusion or error matrix generated by the Random Forest algorithm, showing the success rates in assigning a neuronal type to each neuron. Colour legend correspond with the accuracy of the prediction, 1 being maximum accuracy.

### Supplementary Tables Legends

**Supplementary Table 1. Synaptic hippocampal proteomes characterised in this study.**

**Sheet #1:** Proteins identified by our proteomics workflow from a biochemical preparation of postsynaptic density fractions from total hippocampus.

**Sheet #2:** Reference synaptic proteome used in this study. Produced by combining proteins in sheet #1 with those previously identified in the PSDII fraction published by Distler et al[1].

**Sheet #3:** List of all proteins identified by the Scaffold software from MS/MS data in synaptic fractions from the three hippocampal layers. Proteins common with the reference proteome are indicated. Proteins identified by the Progenesis software with at least two unique peptides are also indicated.

**Sheet #4:** Proteins identified by scaffold in only one of the three synaptic types studied.

**Supplementary Table 2. Analysis of protein abundance and differential protein expression between synaptic types.**

**Sheet#1:** Protein abundance data generated by MSqROB[2,3] from peptide abundance data.

**Sheet#2:** Left of black bar, Proteins with statistically highest expression in one synaptic type. Statistics, one-way ANOVA. FDR correction for multiple testing was performed. Log2 of Fold Chance (FC) and corrected p-values (q-value) are provided. Right of black bar, Protein abundance differences between pairs of synaptic types. Statistics, Student's T-test. FDR correction for multiple testing was performed. Log2 of Fold Chance (FC) and corrected p-values (q-value) are provided.

**Supplementary Table 3. Comparative analysis of protein and RNA expression data from proteins differentially expressed in synapses from the trisynaptic loop.**

**Sheet#1:** Allen Brain Atlas (ABA) *in situ* hybridization (ISH) data was manually inspected for each of the 283 proteins showing differential expression in one of the synapses from the trisynaptic loop (columns E to G). We determined in how many of the 4 brain regions forming the synapses from the trisynaptic loop (Entorhinal cortex Layer II, dDG, CA3 and CA1) was ISH data highest (columns H to K). We compared this information with our proteomics data and stablished if expression levels were concordant or not with protein levels at synapses (column M).

**Sheet#2:** Synaptic genes significantly up or down regulated in excitatory neurons from one of the 4 brain subregions constituting the trisynaptic circuit of the hippocampus: CA1 (dorsal), CA3 (dorsal), dentate gyrus and entorhinal cortex. RNA sequencing data taken from the Allen Brain Cell atlas (ABCA). The ABCA distinguishes dorsal from ventral neurons in the CA1 and CA3 subfields. As the proteomics data was generated from the dorsal hippocampus, we selected dorsal neurons form the ABCA for this analysis. Log2 fold changes and p-values are indicated.

**Supplementary Table 4. Signalling pathways and GO terms significantly overrepresented in proteins with highest expression in one synaptic type.**

**Sheet#1:** The analysis with pathfindR retrieved the following Signalling Pathways as significantly overrepresented amongst protein with highest expression in each synaptic type. Signalling pathways were retrieved from the following databases: Reactome, KEEG and Wikipathways (WP). Fold enrichments are provided, these are calculated as the number of proteins observed in a pathway or term relative to the number expected by chance. The PathfindR metrics occurrence, support, lowest and highest p-values, cluster and status are also provided. Protein with no expression difference between synapses belonging to each pathway are also shown (column K). Proteins from each pathway with highest expression in one synaptic type are indicated (column L).

**Sheet#2:** The analysis with pathfindR retrieved the following GO terms as significantly overrepresented amongst protein with highest expression in each synaptic type. GO terms from the following domains were investigated: Molecular Function (GOMF), Biological Process (GOBP) and cellular component (GOCC). Fold enrichments are provided, these are calculated as the number of proteins observed in a pathway or term relative to the number expected by chance. The pathfindR metrics occurrence, support, lowest and highest p-values, cluster and status are provided. Proteins with no expression difference between synapses belonging to each term are shown (Col. K). Proteins from each term with highest expression in one synaptic type are indicated (Col. L).

**Sheet#3:** Summary of pathways and terms identified as 'Representative' for networks (clusters) of proteins with highest expression in different synaptic types, as determined by PathfindR. A representative pathways or term is the one with the lowest p-value amongst those identified for a protein network.

**Supplementary Table 5. Genes coding for synaptic proteins that have differential RNA expression levels between neuronal classes.**

**Sheet#1:** Table with the number of genes found with a statistically significant up- or down-expression in each neuronal class.

**Sheet#2:** List of genes significantly up- or down-regulated in each class. The ratio of genes up vs. down-regulated is also provided.

**Sheet#3:** List of genes significantly up-regulated in one or two classes.

**Supplementary Table 6. Genes coding for synaptic proteins that have differential RNA expression levels between neuronal types of the same class.**

**Sheet#1:** Table with the number of genes found with a statistically significant up- or down-expression between neuronal types of each class. Neuronal type names as previously published.

**Sheets#2, 4, 6, 8,10, 12, 14 and 16:** Lists of genes significantly up- or down-regulated between neuron types of each of the eight classes investigated.

**Sheets#3, 5, 7, 9,11, 13, 15 and 17:** List of genes significantly upregulated in one or two neuronal types within each class.

**Sheet#18:** Summary table of genes coding for synaptic proteins significantly upregulated in one or two neuronal types.

**Supplementary Table 7. Signalling pathways and GO terms enriched among genes upregulated in different neuronal types.**

**Sheet#1:** List of representative pathways from the databases Reactome, KEGG and Wikipathways identified by pathfindR for synaptic genes upregulated in different neuronal types.

**Sheet#2:** List of representative GO terms identified by pathfindR for synaptic genes upregulated in different neuronal types.

**Supplementary Table 8. Top 1000 genes contributing to the transcriptomics-based classification of excitatory neurons and analysis of the signalling pathways and GO terms associated to them.**

**Sheet#1:** List of the 1000 proteins mostly contributing to the classification of excitatory neurons into classes, as determined by the Random Forest method.

**Sheet#2:** List of the 1000 proteins mostly contributing to the classification of excitatory neurons into types, as determined by the Random Forest method.

**Sheet#3:** Representative terms (Signalling Pathways and GO terms) identified by Pathfinder from the synaptic genes among the top 1000 most contributing to the classification of excitatory neurons.

**Sheet#4:** Representative terms (Signalling Pathways and GO terms) identified by Pathfinder from the non-synaptic genes among the top 1000 most contributing to the classification of excitatory neurons.

**Sheet#5:** Summary of the synaptic and non-synaptic terms used in Figure 6g and h.

**Supplementary Video. Manual dissection of hippocampal subfields.**

**Source Data:**

Source_Data_1_Iteration_Classes.R: R script to iterate the statistical analysis performed with Seurat to identify genes differentially expressed between neuronal classes.

Source_Data_2_Iteration_Types.R: R script to iterate the statistical analysis performed with Seurat to identify genes differentially expressed between neuronal types.

Source_Data_3_Analysis_Classes.R: R script to generate data tables and graphs for genes differentially expressed between neuronal classes. This script also includes a quality control test to validate differentially expressed genes.

Source_Data_4_Analysis_Types.R: R script to generate data tables and graphs for genes differentially expressed between neuronal Types. This script also includes a quality control test to validate differentially expressed genes.

Source_Data_5_Split_Types.R: R script to obtained data from a subset of neuronal types from the entire transcriptomic database provided by the ABCA.

Source_Data_6_pathfindR_Proteomics.Rmd: R script to perform the pathfinder analysis and to generate the heatmaps from the proteomics data.

Source_Data_7_pathfindR_Classes.R: R script to perform the pathfinder analysis and to generate the heatmaps from transcriptomics data of neuronal classes (ABCA).

Source_Data_8_pathfindR_Types.R: R script to perform the pathfinder analysis and to generate the heatmaps from transcriptomics data of neuronal types (ABCA).

Source_Data_9_ Random_Forest.ipynb: Python code to perform the Random Forest analysis on transcriptomic data from the ABCA.

**Data and code Availability**

All the data generated by the bioinformatics analysis performed in this manuscript can be found in the supplementary tables.

Mass spectrometry proteomics data has been deposited to the ProteomeXchange Consortium via the PRIDE partner repository[4] with the dataset identifiers PXD052901 and PXD052913.

All custom-made code is available from GitHub:

https://github.com/Alex-Bayes/Synaptic-Proteome-Diversity

**Acknowledgments**

RRV, DdCB, OZR and AB financial support was provided by: PID2021-124411OB-I00 and RTI2018-097037-B-I00 (MINECO/MCI/AEI/FEDER, EU), Award AC17/00005 by ISCIII through AES2017 and within the NEURON framework, Ramón y Cajal Fellowship (RYC-2011-08391p), IEDI-2017-00822, Universitat Autònoma de Barcelona and AGAUR (2017 SGR 1776 and 2021 SGR 01005). DdCB thanks AGAUR/Generalitat de Catalunya/FEDER, EU for 'Ajuts per a la contractació de personal investigador novel (FI)' Ref.2020FI_B00130. All authors thank the CERCA Programme/Generalitat de Catalunya for institutional support. JP, AC and DS were supported by grant from Ministerio de Universidades, Ciencia y Innovación Innovación (PID2020-119932GB-I00; MCIN/ AEI /10.13039/5011000011033) and the María de Maeztu (MDM-2017-0729 to Institut de Neurociencies, Universitat de Barcelona).

**Author Contributions**

RRV, DdCB, ABP, DAA, OZ, JP, AC and DRV performed experiments. NR, DS, CS and AB designed and supervised all experiments and secured funding. AB and CS wrote the manuscript. All authors reviewed and approved the manuscript.

**Competing interests:**

Authors declare no competing interests.
